# Ca^2+^-driven nanodomain enrichment and plasma membrane proteome remodelling enable bacterial outer membrane vesicle perception in rice

**DOI:** 10.1101/2025.09.17.676730

**Authors:** Ishani Mondal, Hrimeeka Das, Smrutisanjita Behera

## Abstract

1. *Xanthomonas oryzae pv. oryzae* (*Xoo*) releases OMVs; however, the role of *Xoo*-OMVs in *Xoo*-rice interaction has not yet been studied. We investigated the early signalling events underlying *Xoo*-OMV perception by rice to gain insight into early plant-pathogen interactions.
2. We observed that *Xoo*-OMVs are perceived by rice through a rapid and unique Ca^2+^ signal that is essential to evoke an immune response. By confocal imaging and proteomic analysis of the rice PM, we found that the Ca^2+^ signal is required for *Xoo*-OMV-induced nanodomain enrichment and for the aggregation of defence response-associated proteins in rice PM.
3. The differential assembly of proteins in the PM helps the plant defend itself against *Xoo*. In the absence of the early Ca^2+^ signal, *Xoo*-OMVs are not perceived by the plant, resulting in a compromised immune response.
4. Collectively, our study shows that *Xoo*-OMVs are recognised by rice via Ca^2+^ signal-induced nanodomain assembly leading to protein reorganisation in the PM to prepare the plant for an imminent pathogen invasion.

## Introduction

*Xanthomonas oryzae pv. oryzae* (*Xoo*) is a gram-negative bacterium that causes Bacterial Blight of rice (Oryza sativa), leading to a significant loss of rice yield (Niño-Liu et al., 2006). To protect rice against *Xoo*, more than 45 resistant (R) genes have been identified, which are expressed in susceptible varieties through breeding or transgenic approaches to confer resistance against *Xoo*. However, majorly cultivated rice varieties still remain susceptible as it is extremely difficult and tedious to replace hundreds of cultivated varieties by transgenic or gene-edited varieties. To provide sustainable resistance against *Xoo*, a basic understanding of early *Xoo*-rice interaction is necessary. In this context, we explored the inter-kingdom communication between rice and *Xoo* through *Xoo*-released Outer Membrane Vesicles (OMVs). OMVs are spherical nano-proteo-liposomal vesicles released by Gram-negative bacteria to carry out functions like biofilm formation, quorum sensing, virulence, communication, etc., thereby helping the bacteria to adapt to its surroundings (Kulp and Kuehn, 2010; McBroom and Kuehn, 2007; Sartorio et al., 2021).

The role of bacterial OMVs in plant-pathogen interaction is not as extensively explored as it has been in animal-pathogen interaction. Isolation and proteomic characterisation of OMVs from phytopathogenic bacteria, like *Xylella fastidiosa, Xanthomonas campestris pv. campestris* (*Xcc*), *Pseudomonas syringae pv*. *tomato (Pst)*, *Pectobacterium brasiliense,* and *Candidatus liberibacter* (Chowdhury and Jagannadham, 2013; Huang et al., 2022; Maphosa and Moleleki, 2021; Mendes et al., 2016; Sidhu et al., 2008) have identified several OMV-associated proteins, like outer membrane proteins (OMPs), and virulence factors, including T3SS secretion proteins, cell wall-degrading enzymes, porins, lipases, etc. (Chowdhury and Jagannadham, 2013; Jang et al., 2014; Sidhu et al., 2008). OMVs produced by *Xcc* carry MAMPs, such as flg22, EF-Tu, etc. and elicit a PRR-mediated immune response in Arabidopsis (Bahar et al., 2016; Feitosa-Junior et al., 2019; Chalupowicz et al., 2023). *Xcc-*OMVs potentiate nanodomain enrichment, leading to insertion of *Xcc*-OMVs into the host plasma membrane (PM), which further increases the membrane order and reinforces a positive feedback loop to amplify immune responses against the pathogen (Tran et al., 2022). Nanodomains are nanoscale proteo-lipid congregations in the PM that mediate plant responses to biotic and abiotic stress (Gronnier et al., 2017; Jaillais and Ott, 2020; Nagano et al., 2016). For example, specialised nanodomain aggregation is crucial for defining the spatiotemporal specificity of the FLS2 and BRI1-mediated immune pathways in Arabidopsis (Ott, 2017; Bücherl et al., 2017). However, what drives such signal-specific nanodomain enrichment remained unknown.

Parallel to nanodomain enrichment, Ca^2+^ signalling is another indispensable early event for a plant’s immune response. For example, sustained Ca^2+^ increase is observed during the HR response in Arabidopsis challenged by *Pseudomonas syringae* (Grant et al., 2000). Ca^2+^ permeable channels like CNGC2 and CNGC4 are important in shaping PTI-induced Ca^2+^ signals (Ali et al., 2007; Tian et al., 2019). MAMPs like flg22, chitin, elf 18, and pep1, can induce stimulus-specific Ca^2+^ signatures in Arabidopsis mediated by RLCKs, PBL1, and BIK1 (Cao et al., 2017; Keinath et al., 2015; Ranf et al., 2014, 2015). Interestingly, CPK3 can phosphorylate the intrinsically disordered domain (IDD) of REM1.3, a PM nanodomain-associated protein, and thereby influence the function of REM1.3 during plant-virus interaction (Perraki et al., 2018), indicating the role of Ca^2+^-binding proteins in nanodomain clustering. These findings prompted us to investigate the involvement of Ca^2+^ signals in nanodomain aggregation and signal transduction during perception of OMVs by plants.

In this work, we have explored the early events during *Xoo*-rice interaction. We have isolated OMVs from *Xoo,* identified the *Xoo-*OMV*-*associated proteins and examined how rice (*Oryza sativa subsp. indica*) responds to *Xoo-*OMVs. To this end, we have used one susceptible variety of rice, Khitish, and a resistant variety of rice, CR 800. CR 800 expresses three R genes- *xa13*, *xa5,* and *Xa21,* that render it resistant to *Xoo* infection (Mohapatra et al., 2023). Through proteomic analysis, we have found that *Xoo-*OMVs carry abundant MAMPs like EF-Tu, flagellin, Ax21, etc. and elicit a rapid and unique Ca^2+^ signal in the roots of both host rice and non-host Arabidopsis. We have further witnessed that *Xoo-*OMVs can rapidly increase the lipid order of the rice PM and facilitate sterol-mediated insertion of the *Xoo-*OMVs into the PM. We have found that both the above events are Ca^2+^-dependent. Furthermore, the immune response of CR 800 against the *Xoo*-OMVs is diminished in Ca^2+^-deprived condition. Quantitative proteomic analysis of the *Xoo*-OMV-treated CR 800 PM revealed that *Xoo*-OMVs increase the abundance of defence response-related proteins in the PM in the presence of Ca^2+^ but not in a Ca^2+^-chelated condition. Collectively, our results describe the Ca^2+^-dependent relay of events during the perception of OMVs by the host plant and contribute to the understanding of early host-pathogen interaction.

## Materials and Methods

### Plant and bacterial samples

Khitish (susceptible to *Xoo,* does not have R genes) and CR 800 (resistant to *Xoo*, contains R genes, Xa21, xa3, xa15) (*Oryza sativa subsp. indica*) were grown hydroponically at 30 °C under a 14-hour light / 10-hour dark condition, with 70-80% humidity. Col-0-RGECO1 (Col-0 plants expressing the genetically encoded Ca^2+^ indicator (GECI), RGECO1 were grown in 0.5 strength MS media at 22 °C. *Xanthomonas oryzae pv. oryzae* strain BXO43 (rifampicin derivative of BXO1, a Bengal isolate, (Rajagopal et al., 1997) was grown in peptone sucrose (PS) media (O.D._600_ of 0.6) with rifampicin (50 µg/ml) at 28 °C.

### Isolation of outer membrane vesicles

*Xoo*-OMVs were isolated and purified according to the protocol of Mordukhovich and Bahar, 2017. Briefly, the supernatant of the BXO43 culture was filtered through a 0.45 μm syringe filter to obtain the cell-free supernatant. It was ultracentrifuged at a speed of 100000g at 4 °C for 2 hours and filtered again through a 0.22 μm filter followed by another wash to remove contaminant proteins. This crude *Xoo-*OMV preparation was loaded on top of a density gradient of 45%, 40%, 35%, 30%, 25%, and 20% of OptiPrep diluent buffer (Sigma) for purification. Each fraction was collected and washed twice to remove the diluent buffer. The resultant purified *Xoo-*OMV pellet was weighed and re-suspended in sterile deionized water or in Ca^2+^ imaging buffer as per the requirement of the experiment, at a concentration of 100 mg/ml fresh weight. Isolated *Xoo-*OMVs were stored at 4 °C and utilized within 3 days of isolation. The total protein content of the *Xoo-*OMVs was quantified using Bradford assay (Biorad).

### Nanoparticle Tracking Analysis (NTA)

The *Xoo-*OMVs were diluted 200 times in sterile water and injected into the laser chamber of Nanosight NS300. The videos were analysed and plotted in a graph using the NTA 3.2 Dev Build 3.2.16 software.

### Atomic force microscopy (AFM)

*Xoo* cells were diluted with PBS to an O.D. of 0.2. The diluted *Xoo* cells were incubated at 28 °C for 1 hour. 3 μl of the cells were dropped on a freshly cleaved mica sheet. The sheet was dried inside a vacuum desiccator for 1 hour and visualized with the help of the AFM-MFP-3D (Oxford Instruments) in tapping mode.

### Transmission electron microscopy (TEM)

*Xoo-*OMVs were fixed in 2% glutaraldehyde, and 5 μl of the sample was dropped on a copper grid. The sample was stained with 1% uranyl acetate and visualized under a Jeol-JEM-2100 Plus transmission electron microscope.

### Isolation of PM-associated proteins from CR 800

7-day-old CR 800 seedlings were incubated for 1 hour in sterile water (Control) or in *Xoo*-OMVs (OMV). For Ca^2+^-chelation, seedlings were treated with 2 mM EGTA for 30 minutes before incubation in *Xoo*-OMVs (EGTA-OMV) for 1 hour. PM was isolated from the roots according to the protocol described in (Uemura et al., 1995). Briefly, roots of ∼ 4000 seedlings were homogenised in homogenising buffer (0.5 M sorbitol, 50 mM Mops-KOH (pH 7.6), 5 mM EGTA (pH 8.0), 5 mM EDTA (pH 8.0), 5% (w/v) PVP-40, 0.5% (w/v) BSA, 2.5 mM PMSF,4 mM SHAM, 2.5 mM DTT). The homogenate was centrifuged to remove debris and ultracentrifuged to obtain the microsomal fraction. The microsomal fraction was enriched with PM proteins by a repeated two-phase partitioning system using PEG-dextran (6%, w/w). The enriched PM proteins were washed twice and re-suspended in PM suspension buffer (10 mM Mops-KOH (pH 7.3), 2 mM EGTA (pH 8.0), 0.25 M sucrose) for further proteomic analysis. Three such independent biological replicates were performed for each sample (Control, EGTA-OMV, OMV).

### SDS-PAGE and in-gel trypsin digestion

Proteins were precipitated from the *Xoo-*OMV samples using water: methanol: chloroform=4:3:1. The protein pellet was recovered from the interface of the aqueous and organic phases, washed with methanol and dried. CR 800-PM proteins and *Xoo*-OMV proteins were resuspended in 1X Tris-glycine sample loading buffer (Invitrogen). 50 μg of each replicate of the respective protein samples were run separately in a 12% SDS-PAGE gel. After brief staining with Coomassie blue G-250 the protein bands were cut, de-stained with ammonium bicarbonate and dehydrated with ACN. The proteins were digested with trypsin for 15 minutes, followed by protein extraction for 24 hours. An additional step of 1 hour of protein extraction was carried out in order to maximize the peptide concentration. The peptides were lyophilized and stored at −20 °C until further analysis through the ESI-MS/MS system.

### LC-ESI-MS/MS of digested peptides

The digested peptides (of both *Xoo*-OMV proteins and CR 800-PM proteins) were reconstituted in 10 μl of 1% formic acid and 0.75 μg of peptides were subjected to the nano-flow (75 μm x 500 mm, EASY-Spray™ PepMap™ Neo C18 column, with 100 Å pore size) of an Orbitrap Fusion Lumos mass spectrometer (Thermo Fisher Scientific). The system was run for 140 minutes with Solvent A=0.1% formic acid and Solvent B=80% ACN 0.1% formic acid. The peptides were eluted at a constant flow of 0.3 μl/min, starting with 5% solvent B to 75% solvent B up to 125 minutes, then decreasing to 5% solvent B until 140 minutes. The scan was performed in a data-dependent mode with 20 dependent scans with a resolution of 60000 and scan range of m/z from 350-2000. The peaks of the significant proteins were plotted directly using X-calibur software™ (Thermo Fisher Scientific).

### Protein identification, quantitation, and GO enrichment analysis

The *Xoo*-OMV proteins were identified using Proteome Discoverer software with Sequest HT and Percolator. The proteins were identified. The identification was performed with: enzyme: trypsin; missed cleavage: 2; precursor mass tolerance: 10 ppm; peptide charge: +1, +2 and +3; ion type: monoisotopic against the reference proteome of *Xoo*, strain KACC10331/KXO85, Taxon ID. 291331, of the UniProt database. Functional annotations of *Xoo*-OMV-associated proteins were designated using String Database (https://string-db.org/).

CR 800-PM proteins were identified, and the grouped abundances, abundance ratios, and p-values of the proteins in the respective samples were quantified through label-free quantification (LFQ) technique, using the consensus workflow, CWF_Comprehensive_Enhanced Annotation_LFQ_and_Precursor_Quan and processing workflow as PWF_Tribrid_Precursor_Quan_and_LFQ_ITHCD_CID_SequestHT_Percolator. The biological, molecular functions, subcellular localisations and KEGG pathways of differentially abundant proteins (DAPs) (≥1.5-fold change, p<0.05) were identified using String Database (https://string-db.org/). Their enrichment is depicted in a lollipop graph, showing strength (Log10(observed/expected)) and FDR (corrected p-values). The grouped abundance of important proteins was shown in heat maps with ChiPlot (https://chiplot.online/).

### Ca^2+^ imaging of *Arabidopsis thaliana* Col-0 RGECO1 seedlings

4-day-old Col-0-RGECO1 seedlings were placed carefully inside a chambered slide with an agar block on top and 10 μl of Ca^2+^ imaging buffer (5 mM KCl, 10 mM MES, and 10 mM CaCl_2_, pH 5.8, adjusted with TRIS) (Behera et al., 2015) was added to keep the seedlings moist during imaging. 20 μl of *Xoo*-OMVs was added inside the chamber after 5 minutes of imaging. The seedlings were visualized under a HC PL APO CS 10X/0.40 dry objective of Leica SP8 confocal microscope. The seedlings were excited at 561 nm to collect emission between 605 to 665 nm. The roots were imaged at an interval of 10 seconds for 30 minutes. The images were analysed using ImageJ software and fractional fluorescence change, ΔF/F, was calculated according to (Keinath et al., 2015). Briefly, the fluorescence intensity of each frame was background corrected and normalized with the average intensity of the baseline, i.e., the first five minutes as (F-F_0_)/F_0_. Three regions of interest (ROIs) were selected, ROI1- meristematic region of the root; ROI2- distal elongation region of the root; ROI3- central elongation region of the root. GraphPad Prism was used to draw the line curve representing the spatiotemporal change in Ca^2+^ levels at individual time points (n = 6).

### Calcium Green 1, AM staining of rice seedlings

7-day-old Khitish and CR 800 seedlings were stained with Calcium Green1, AM, (Invitrogen) (15 μM) in presence of 0.5 mg/ml phytostigmine salicylate (Sigma) for better dye loading. The stained rice roots were kept inside chambered slides and covered by an agar block. 75 μl sterile water was added to keep the roots moist while imaging and visualized under a Zeiss LSM800 confocal laser scanning microscope. The roots were excited at 488 nm at an interval of 1 minute under a PLAN APO 10X dry objective with a 0.5X magnification, and emission was collected in the range of 500-600 nm. After 5 minutes of imaging, 150 μl *Xoo*-OMVs was added to the chambered slide. The background-corrected and normalised fluorescence intensity values (ΔF/F) of the epidermis region of the root elongation zone were analysed using the ImageJ Fiji software. The ΔF/F values of seedlings treated with *Xoo*-OMVs or sterile water, respectively, at an interval of 5 minutes were plotted in a histogram using GraphPad Prism software. 6 independent experiments were compared using Student’s t-test.

### qRT-PCR

7-day-old Khitish and CR 800 seedlings were incubated in *Xoo-*OMVs for 1 hour, followed by incubation in sterile water for 1 hour (OMV) or 2 hours in sterile water (control) or pre-treated with calcium chelators, BAPTA-AM (25 μM) or EGTA (2 mM) for 30 minutes, followed by incubation in *Xoo-*OMVs for 1 hour then in sterile water for 1 hour. RNA was isolated from the root tissue of the rice seedlings, with subsequent cDNA preparation. qRT PCR was performed to quantify the expression of *PR10b* gene with the following primers: qActin-F (5′ ATCCTTGTATGCTAGCGGTCGA-3′), qActin-R (5′-ATCCAACCGGAGGATAGCATG-3′); PR10b-F (5′-TGTGGAAGGTCTGCTTGGAC-3′), PR10b-R (5′- CCTTTAGCACGTGAGTTGCG-3′) (Liu et al., 2020). Three independent replicates of the experiment had been performed with each replicate consisting of 5 seedlings.

### Quantification of PM lipid order

7-day-old Khitish and CR 800 roots and (leaf) protoplasts, collected from the leaf tissues, were treated with *Xoo-*OMVs (25 mg/ml) for 30 minutes, 60 minutes, or 90 minutes. The roots and protoplasts were pre-treated with BAPTA-AM (25 μM), or EGTA (2 mM), or MβCD (2 mM) (sterol-depleting agent), respectively, for 30 minutes followed by incubation in *Xoo*-OMVs as mentioned. The roots and protoplasts were stained with 4 μM Di-4-ANEPPDHQ for 10 minutes and viewed under a Leica SP8 confocal HC PL APO CS 60X oil immersion objective, excited at 488 nm, and emission was collected at two different ranges from 500-580 nm (ordered phase) and 620-750 nm (disordered phase). The GP values (+1 to −1) were calculated according to the protocol of Owen et al., 2012, (n≥6).

### Fusion assay of OMV with FM4-64

*Xoo-*OMVs were stained with 1.5 mM FM4-64 for 10 minutes on ice in the dark. Stained *Xoo-*OMVs were washed to remove any residual stain. The stained *Xoo*-OMV (FM4-64*-*OMV) pellet was resuspended in sterile water to a concentration of 25 mg/ml, kept at 4 °C, and used within 2 days. Sterile water subjected to similar staining and washing steps was used as mock treatment (control). 7-day-old Khitish and CR 800 roots and leaf protoplasts were incubated with FM4-64-OMVs or the mock treatment for 30 minutes, 60 minutes, or 90 minutes, respectively. The seedlings were imaged under a HC PL APO CS 60X oil immersion objective of Leica SP8 STED confocal microscope and excited at 568 nm, with emission collected from 700-795 nm. The seedlings were pre-treated with BAPTA-AM (25 μM) or EGTA (2 mM), or MβCD (2 mM) for 30 minutes when mentioned, (n≥7)

### Statistical analysis

GP values and fluorescence intensity values of Fig. 3 and 4, respectively, were calculated using 15 ROIs from each image corresponding to individual seedlings with the help of ImageJ Fiji software. The graphs, including the histograms, box plots, and line curves, were plotted to visualize these values through GraphPad Prism software. One-way ANOVA was used to deduce the statistical difference between the samples using GraphPad Prism software unless specified. The p-value was taken as 0.05 with ns, not significant, *P≤0.05; **P≤0.01, ***P≤0.001, ****P≤0.0001.

**Fig. 1.**
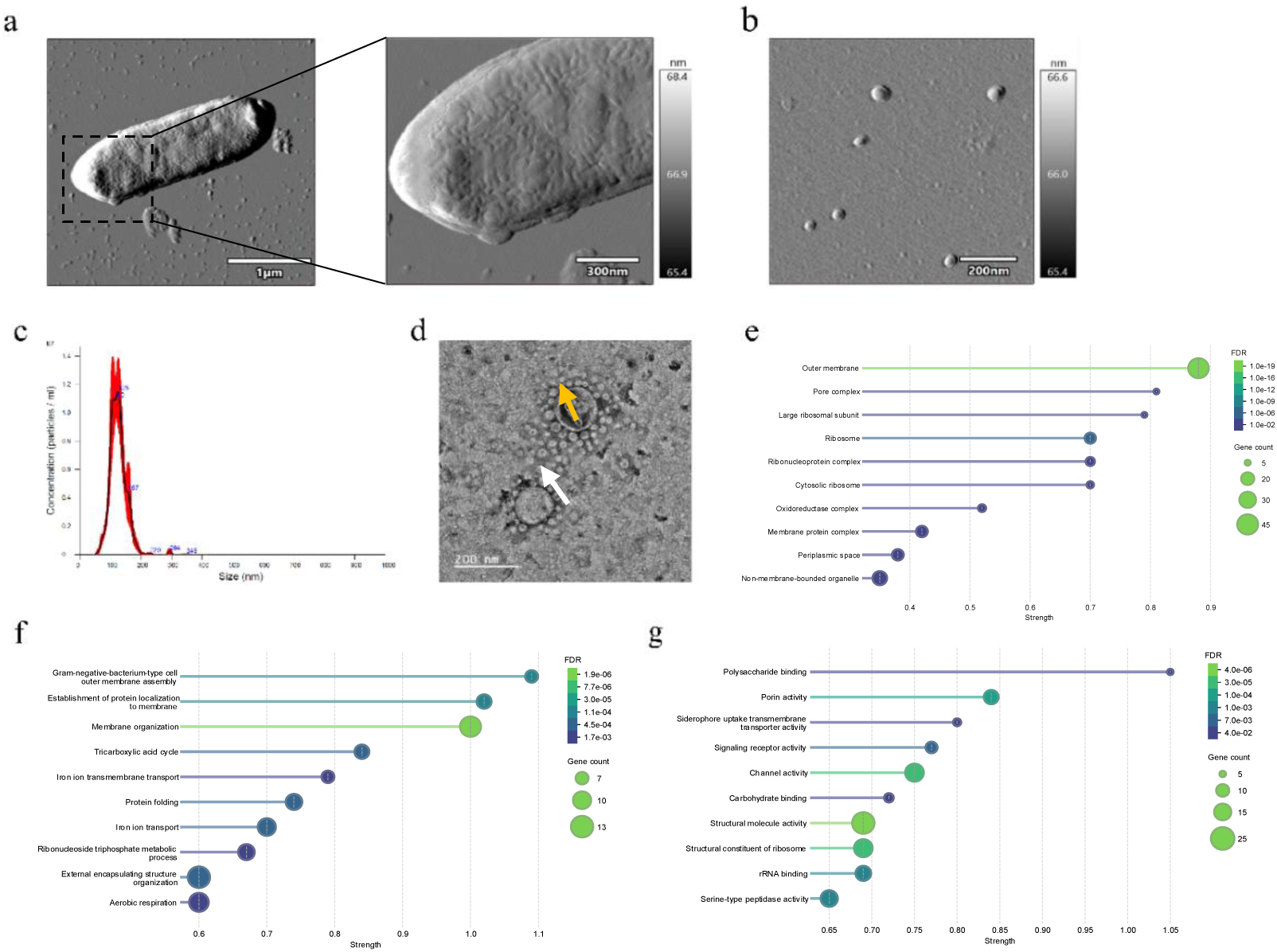
Characterization of *Xoo-*OMVs and identification of their protein content. a, AFM images showing the release of BXO43 OMVs into their surroundings by blebbing. b, AFM image of spherical *Xoo*-OMVs. c, the hydrodynamic size range of the *Xoo-*OMVs (125.8 +/-1.6 nm) exhibited by NTA. d, TEM image of *Xoo-*OMVs including smaller vesicles (10-60nm), indicated by the yellow arrow, and bigger vesicles (100-150nm), indicated by the white arrow. e, Subcellular localization of 423 proteins identified by proteomic analysis of *Xoo-*OMVs (n=4) are designated using String Database. f and g, predicted biological, and molecular functions of the proteins, appointed using String Database are represented in lollipop charts, with strength (log_10_(Expected/Observed)) in the x-axis.

**Fig. 2.**
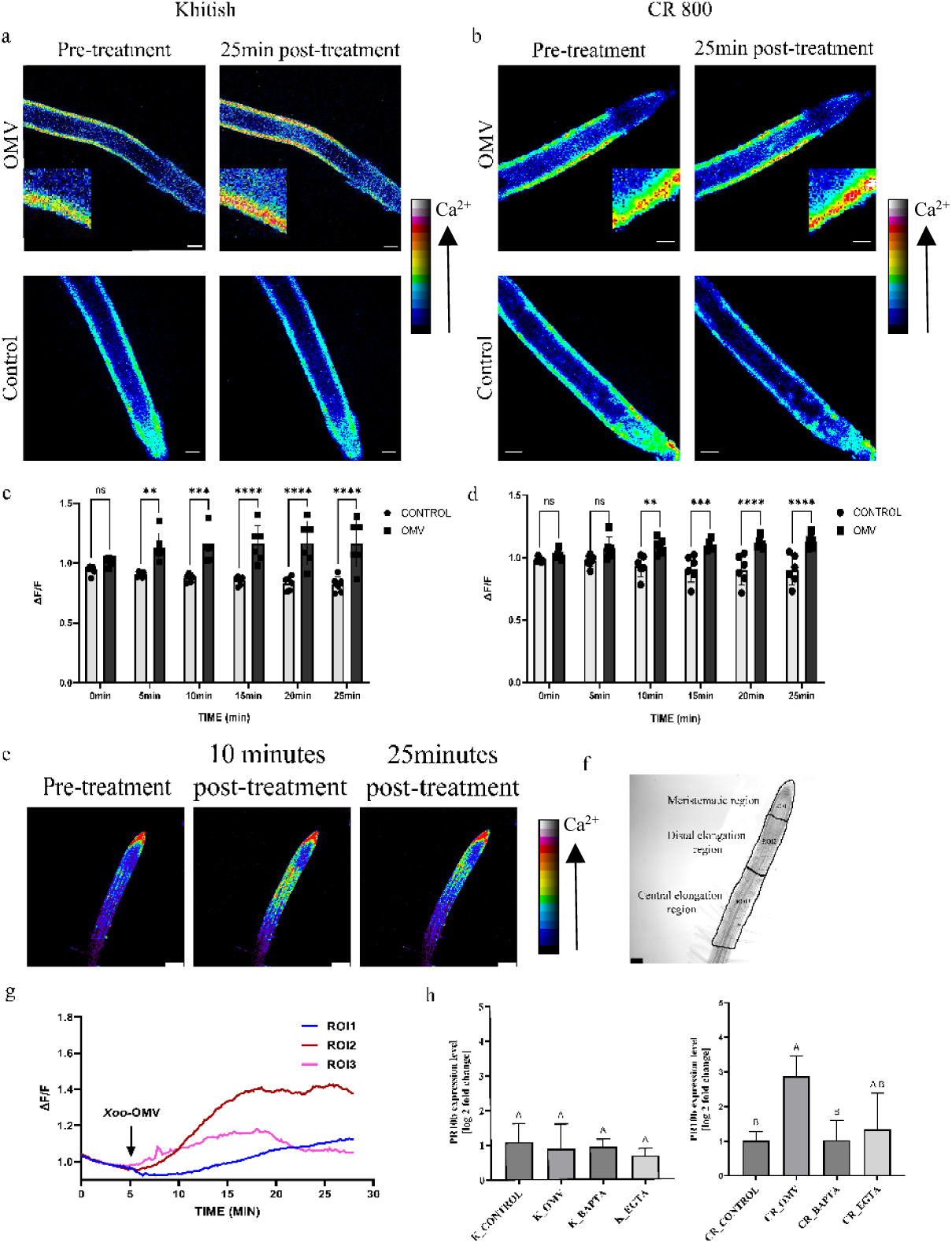
*Xoo*-OMVs induce cellular Ca^2+^ elevation in rice and Arabidopsis roots. a and b, 7-day-old Khitish and CR 800 roots, respectively, stained with Calcium Green1, AM, showing an increase in cellular Ca^2+^ when treated with *Xoo-*OMVs. No Ca^2+^ elevation was observed with solvent treatment, scale bar=125 μm. c and d, the relative fluorescence of the Khitish and CR 800 roots after treatment with *Xoo-*OMVs and solvent control were plotted as a histogram at intervals of 5 minutes using GraphPad Prism software (n=6). Student’s t-test was done to deduce the statistical difference between the fluorescence of control and *Xoo-*OMV-treated roots at each time point, n=6. e, 4-day-old Arabidopsis Col-0 roots, expressing Ca^2+^ indicator protein RGECO1 treated with *Xoo-*OMVs. showing an increase in fluorescence, indicating a Ca^2+^ spike after treatment with *Xoo-*OMVs, scale bar=50 μm. f, selection of ROIs, namely, the meristematic region (ROI1), the distal elongation region (ROI2), and the central elongation region (ROI3). g, the average line curve (n=5) showing the Ca^2+^ signal induced by *Xoo-*OMVs in different ROIs of the Arabidopsis roots with respect to time. The black arrow indicates the time of application of *Xoo-*OMVs. h, the relative expression of the *PR10b* gene after incubation in *Xoo-*OMVs for 1 hour (in the presence and absence of Ca^2+^-chelator) in CR 800 and Khitish roots, (n=3 independent replicates). The statistical differences have been calculated through one-way ANOVA using GraphPad Prism software (ns, not significant, *P≤0.05; **P≤0.01, ***P≤0.001, ****P≤0.0001).

**Fig. 3.**
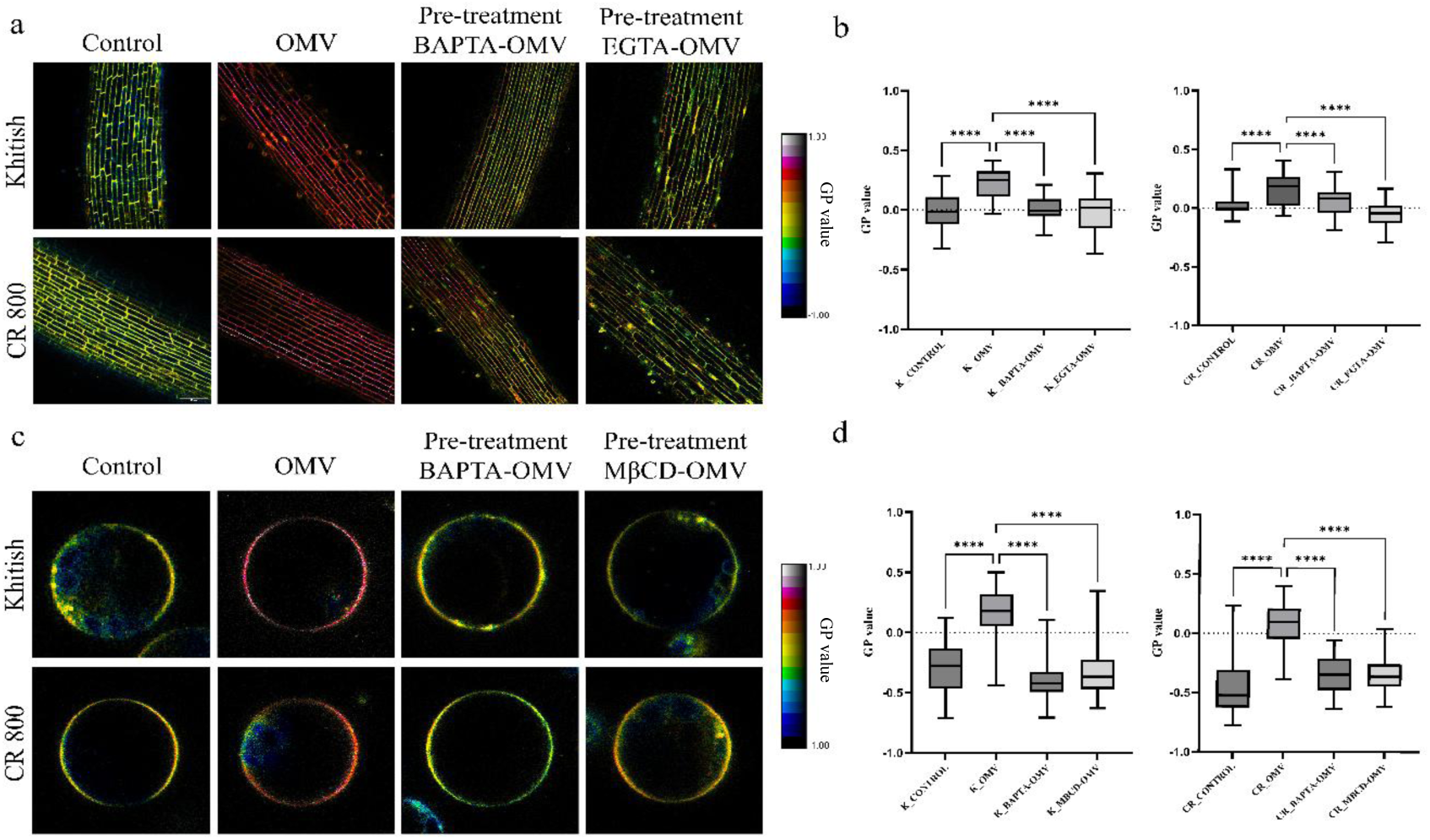
Ca^2+^ and sterol-dependent lipid order increase in rice PM after incubation in *Xoo-*OMVs. a, 7-day-old Khitish and CR 800 roots incubated with *Xoo-*OMVs for 30 min and stained with Di-4-ANEPPDHQ showing an increase in the lipid order of rice PM (n≥6 seedlings). 30-minute pre-treatment with BAPTA-AM (25 μM) or EGTA (2 mM) inhibits lipid order increase. b, GP values (−1 to +1) have been calculated using ImageJ software and plotted in a Box-plot using GraphPad Prism software, the corresponding statistical significance have been calculated through one-way ANOVA (p≤0.05), c, leaf protoplasts isolated from Khitish and CR 800 showing lipid order rise after incubation with *Xoo-*OMVs. BAPTA-AM (25 μM) or MβCD (2 mM) pre-treated protoplasts not showing significant rise in lipid order even after *Xoo-*OMV exposure. d, GP values of the protoplast membrane of Khitish and CR 800, represented in a box plot. The statistical significance has been calculated by one-way ANOVA using GraphPad Prism software (ns, not significant, *P≤0.05; **P≤0.01, ***P≤0.001, ****P≤0.0001).

**Fig. 4.**
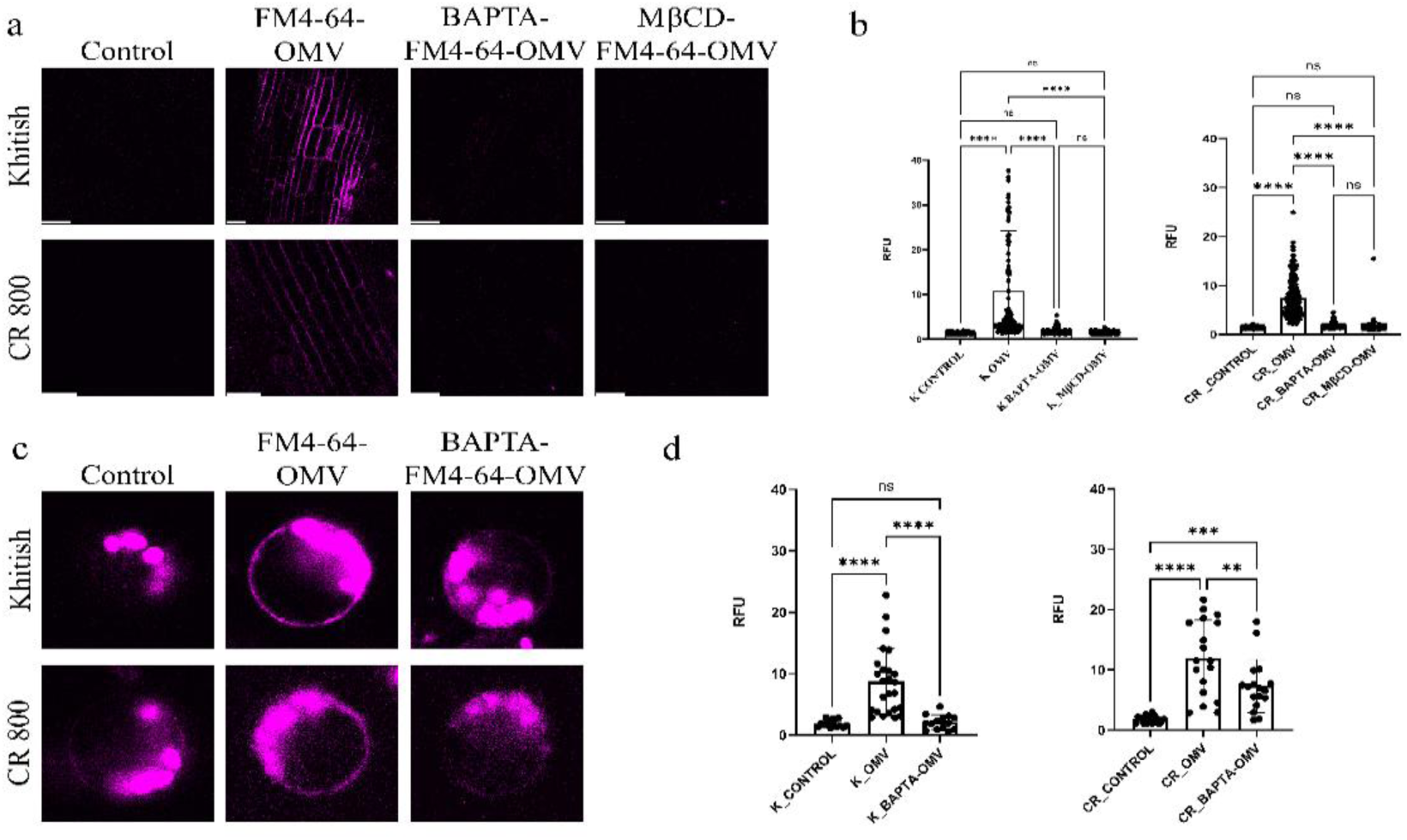
*Xoo-*OMVs get inserted into rice PM. a, 7-day-old Khitish and CR 800 roots showing a significant decrease in *Xoo-*OMV insertion on Ca^2+^-chelation and sterol-depletion with BAPTA-AM (25 μM) and MβCD (2 mM), respectively (n≥7 seedlings), scale bar=20 μm. b, histogram showing the statistical difference between the relative fluorescence units of each sample, calculated using one-way ANOVA by GraphPad Prism. c, FM4-64 stained *Xoo-*OMVs inserted into the PM of leaf protoplasts with and without pre-treatment with BAPTA-AM (25 μM). d, GraphPad Prism was used to create the histograms and analyse the statistical difference between the protoplast samples through one-way ANOVA, (ns, not significant, *P≤0.05; **P≤0.01, ***P≤0.001, ****P≤0.0001).

## Results

### *Xoo-*OMV*-*associated proteins include MAMPs and virulence-associated proteins

We isolated OMVs from the BXO43 strain of *Xoo*. Through Atomic Force Microscopy (AFM) we observed that *Xoo* spontaneously releases spherical OMVs into the surroundings by blebbing (Fig. 1a, b). We determined the hydrodynamic range of the *Xoo*-OMVs using nanoparticle tracking analysis (NTA) to be 125.8 ± 1.6 nm (Fig. 1c). Transmission electron microscope (TEM) images of the *Xoo-*OMVs revealed the presence of smaller vesicles (10-60 nm) and larger vesicles (100-150 nm), indicated by a yellow and a white arrow, respectively (Fig. 1d). This corroborates with the previous reported data regarding *Xcc*-OMVs and PXO99A-OMVs, which exhibit a similar size range (Bahar et al., 2016)

We identified 290 proteins through qualitative proteomics of the proteins associated with the purified *Xoo*-OMVs, (Table S1). The identified proteins were grouped according to their functional annotations (biological, molecular functions and subcellular localisations) with the help of String Database. The proteins associated with *Xoo*-OMVs showed maximum enrichment of outer membrane proteins and pore complex proteins involved in outer membrane assembly and organisation of *Xoo* (Fig. 1e, f). Proteins involved in iron-ion transport and siderophore uptake like TonB-dependent receptors (iroN, fyuA, btuB), and iron transporters (fecA, yncD, CirA, etc) are abundant in *Xoo*-OMVs (Fig. 1g). Phosphate acquisition proteins like phoP, pstS, oprO, ppa, fruB in the *Xoo*-OMV proteome suggest the role of OMVs in nutrient acquisition (Table 1).

**Table 1:**
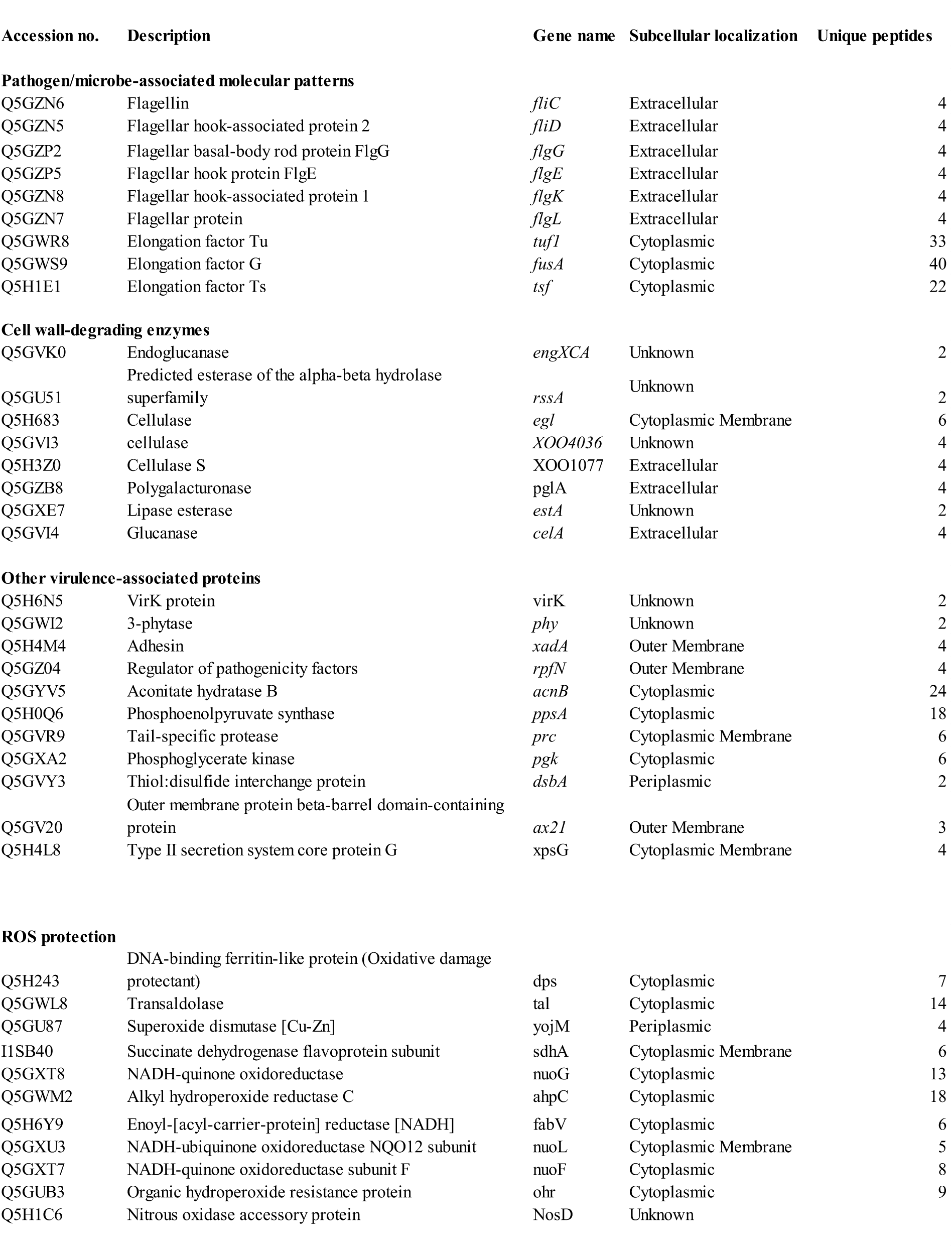
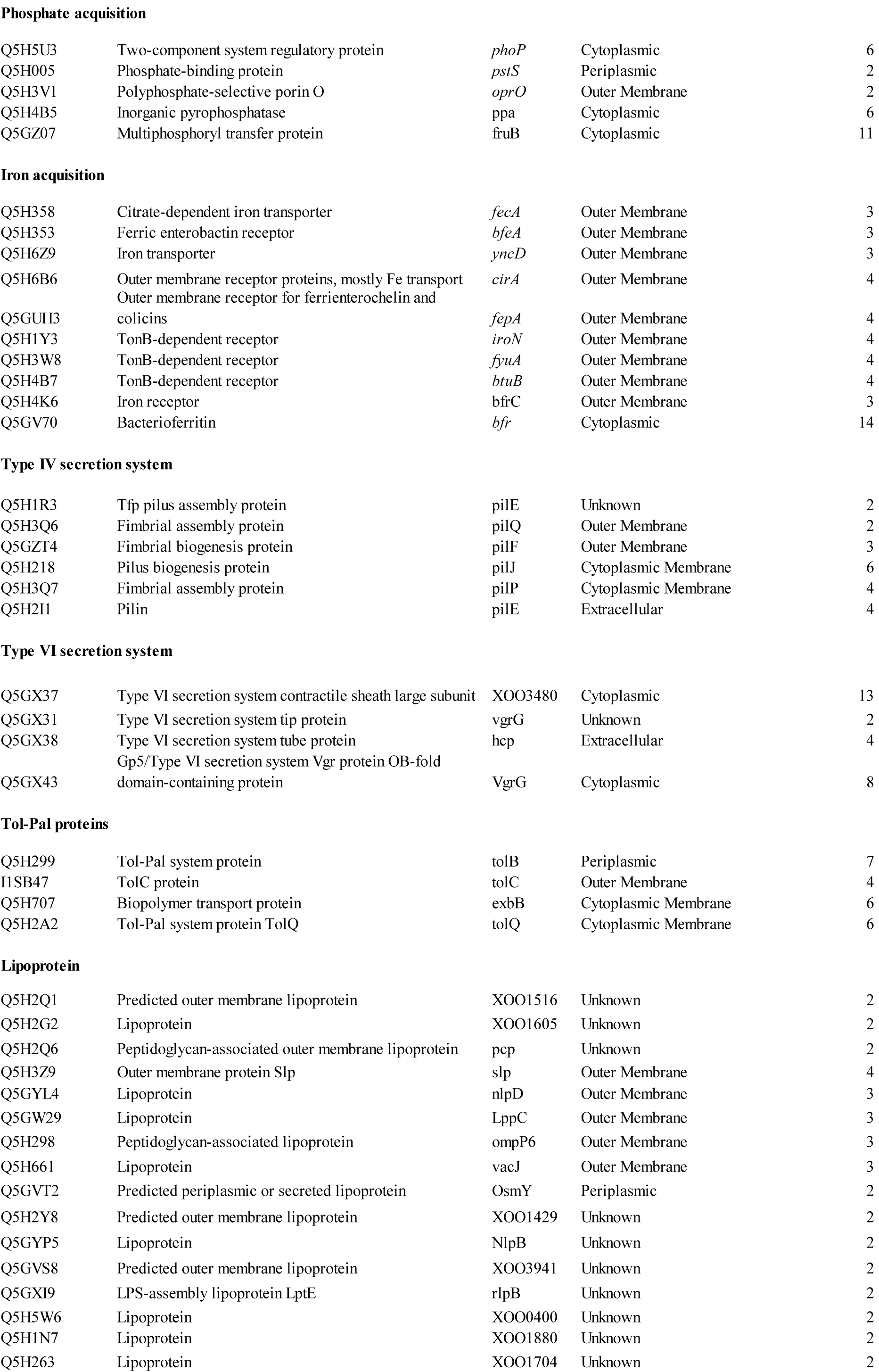
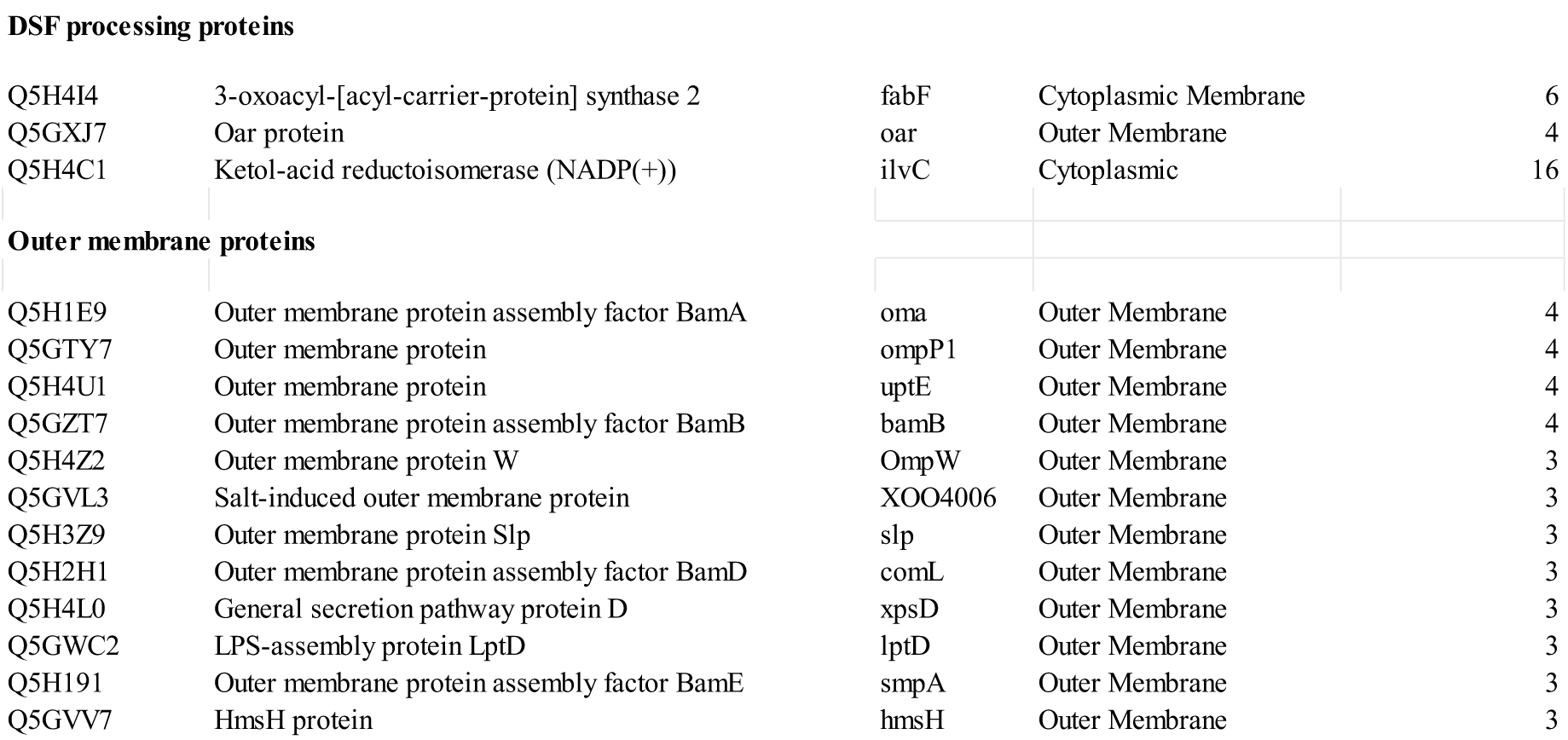
Important proteins identified in Xoo-OMVs.

We report that *Xoo-*OMVs, like *Xcc*-OMVs (Bahar et al., 2016, 2014), carry known PAMPs like Ef-Tu, flagellar proteins (fliC, flgE, flgL, etc), and Ax21. Presence of other virulence-associated proteins like regulator of pathogenicity factors (rpfN), adhesins (xadA), extracellular serine protease (XOO2375), and virulence protein (virK), etc. (Table 1) imply the importance of *Xoo*-OMVs in virulence. xadA facilitates invasion through hydathodes and is crucial for optimal virulence (Das et al., 2009). XOO2375 facilitates colonization through the xylem tissue (Mendes et al., 2016). Presence of several plant-cell-wall-degrading enzymes, including cellulase (egl, XOO4036, XOO1077), endoglucanase (engXCA), esterase (estA), glucanase (celA), xylanase (xynA), etc. and secretory system proteins like T6SS proteins (vgrG, hcp, XOO3480) and T4SS proteins, (including pili-forming subunits, pilE, pilQ, pilF, pilJ, pilP, etc.), suggest the role of *Xoo*-OMVs in *Xoo* infection (Table 1). Moreover, *Xoo-*OMVs carry proteins that facilitate defence against host immune responses (particularly against ROS produced by the plants in response to OMVs), such as transaldolase, yojM, dps, organic hydroperoxide resistance protein (ohr), NADH-quinone oxidoreductase proteins (nuoF, nuoL, nuoG), etc. (Table 1). Overall, analysis of *Xoo*-OMV-associated proteome propose the role of *Xoo-*OMVs in virulence, nutrient acquisition, invasion, colonization, and defence against the host’s immune response.

### *Xoo-*OMVs induce Ca^2+^ signal in both rice and Arabidopsis

Numerous reports describe the involvement of specific Ca^2+^ signals in sensing PAMPs, such as EF-Tu, chitin, and flg22, which induce a Ca^2+^ signal in plant cells (Cao et al., 2017; Keinath et al., 2015; Ranf et al., 2014). The identification of these MAMPs in the *Xoo*-OMV-associated proteome drove us to examine whether *Xoo*-OMVs induce any Ca^2+^ signal in plant cells. To accomplish this, we used a Ca^2+^-indicator stain, Calcium Green1, AM, to stain CR 800 (resistant variety) and Khitish (susceptible variety) seedlings, respectively. Rice leaves are hydrophobic and difficult to stain; as an alternative, we imaged rice roots, as they serve as the entry point for *Xoo* infection through contaminated irrigation water and soil (Naqvi, 2019). We observed an increase in Ca^2+^ concentration (depicted by the false colour code, calculated as ΔF/F) in the epidermal cells of the root distal elongation zone of both Khitish and CR 800 after 10 minutes of *Xoo*-OMV treatment (Fig. 2a, b). No elevation of Ca^2+^ concentration was observed in the roots treated with sterile water. ΔF/F values of each seedling treated with sterile water (control) or *Xoo-*OMVs (OMV) at specific time points were plotted in a histogram for comparison (Fig. 2c, d). Six independent imaging experiments were carried out for each variety.

To further visualize the spatiotemporal propagation of the *Xoo*-OMV-induced Ca^2+^ signals we used transgenic Arabidopsis Col-0 seedlings expressing the genetically encoded Ca^2+^ indicator RGECO1 as the system. We placed 4-day-old Col-0-RGECO1 seedlings in custom-made chambered slides, and performed live-cell imaging as described in the Material and Methods. The seedlings were imaged for 5 minutes in solvent control to obtain a stable baseline, and then 20 µl of 100 mg/ml *Xoo-*OMVs was added to the chamber containing the seedling. We observed a gradual Ca^2+^ signal with an amplitude of 0.5 (ΔF/F) after 10 minutes of purified *Xoo*-OMV application (Fig. 2e, g). To analyse the observed signal, we defined three ROIs around three distinct regions of the root, namely, the meristematic region (ROI1), the distal elongation region (ROI2), and the central elongation region (ROI3) (Fig. 2f). The increase in the Ca^2+^ concentration was most prominently displayed in the distal elongation zone of the root (Fig. 2e, g). In conclusion, *Xoo-*OMVs can induce a Ca^2+^ signal in both the host rice and non-host Arabidopsis within 10 minutes of exposure.

To investigate whether the early Ca^2+^ signal is necessary for the OMV-induced immune response in host plants, we treated 7-day-old Khitish and CR 800 seedlings with *Xoo-*OMVs for 1 hour, with or without pre-treatment with Ca^2+^-chelators (BAPTA or EGTA), as described in the Materials and Methods. We observed that the resistant variety of rice, CR 800, could exhibit an immune response against *Xoo* after treatment with *Xoo*-OMVs, indicated by a three-fold elevation in the expression of *PR10b gene* compared to the incubation in sterile water. On the other hand, Khitish, the susceptible variety of rice, could not evoke any such immune response after exposure to *Xoo-*OMVs (Fig. 2h). Further, the immune response displayed by CR 800 was weakened in a Ca^2+^-deprived condition, indicated by the reduced expression of *PR10b* (Fig. 2h). These results suggest that the early Ca^2+^ signal triggered by *Xoo*-OMVs is crucial for the perception of *Xoo*-OMVs and activation of downstream immune response against *Xoo* in CR 800.

### *Xoo-OMVs* instigate rapid nanodomain enrichment in rice in a Ca^2+^-dependent manner

It has been reported that the fusion of *Xcc*-OMVs with the PM of Arabidopsis leads to an increase in the membrane order (Tran et al., 2022). We inspected whether *Xoo*-OMVs can exert similar effect on rice PM. Furthermore, we investigated the role of Ca^2+^signal generated by *Xoo*-OMVs in changing the membrane order. We used Di-4-ANEPPDHQ, a polarity-sensitive membrane dye (Owen et al., 2012), to assess the state of lipid order of rice PM in the presence of *Xoo-*OMVs. Di-4-ANEPPDHQ shows a spectral shift of 60 nm between disordered and ordered phases of the membrane. The lipid order of the membrane can be quantified using the generalised polarity (GP) values, which range from −1 to +1, indicating the polarity of the membrane and thereby suggesting the order of the membrane. Higher GP values correspond to a highly ordered membrane or a nanodomain-rich membrane (Owen et al., 2012).

We found that after 30 minutes of incubation in *Xoo-*OMVs, the roots of 7-day-old Khitish and CR 800 seedlings showed a significant increase in the GP value of the PM (P≤0.0001, n≥6 seedlings), compared to roots incubated in sterile water (control) (Fig. 3a, b), indicating a highly ordered membrane (Fig. 3a, b). The increased order of the PM was maintained for up to 90 minutes of *Xoo*-OMV treatment (Fig. S1). Conversely, Khitish and CR 800 seedlings pre-treated with EGTA or BAPTA-AM for 30 minutes exhibited a reduced increase in the lipid order of the PM after 30 minutes of incubation. *Xoo-*OMVs (Fig. 3a, b).

We examined whether *Xoo-*OMVs had an analogous effect on the PM of the leaf tissue as well. As it was difficult to stain rice leaves due to their hydrophobic nature, we isolated protoplasts from Khitish and CR 800 leaves and incubated the protoplasts with *Xoo-*OMVs for 30 minutes. The protoplasts stained with Di-4-ANEPPDHQ (3.5 μM) were mounted on a chambered slide for visualisation. The protoplasts incubated with *Xoo-*OMVs displayed significantly higher GP values than the protoplasts incubated with sterile water (control) (Fig. 3c). Similar to the rice roots, leaf protoplasts also displayed a compromised increase in the lipid order of the PM after Ca^2+^-chelation with BAPTA-AM (25 μM) (Fig. 3c, d). In order to verify that the rise of the lipid order of the host membrane is due to nanodomain enrichment, and not merely a biophysical effect, we treated the rice leaf protoplasts with MβCD (2 mM), a sterol-depleting agent. Protoplasts pre-treated with MβCD showed no increase in lipid order upon incubation in *Xoo-*OMVs for 30 minutes (Fig. 3c). The GP values (+1 to −1) of the respective experiments have been plotted in a box plot, and their statistical differences have been calculated using one-way ANOVA with the help of GraphPad Prism software (Fig. 3b, d). Collectively, our observations suggest that the early Ca^2+^ signal is necessary for *Xoo*-OMV-induced nanodomain enrichment in the rice PM.

### Incorporation of *Xoo-*OMVs in rice PM is Ca^2+^-dependent and nanodomain-mediated

OMVs can enter the non-phagocytic host cells through various pathways, especially in a cholesterol-dependent lipid raft-mediated way (Kaparakis-Liaskos and Ferrero, 2015; Schaar et al., 2011). *Xcc-*OMVs get incorporated into the Arabidopsis PM in a sterol-dependent manner (Tran et al., 2022). Accordingly, we investigated the insertion of *Xoo*-OMVs in rice

PM and its Ca^2+^ dependency. Similar to the aforementioned paper, we used FM4-64, a lipophilic dye, to stain the *Xoo-*OMVs, allowing us to track them. *Xoo*-OMVs were stained as described in Materials and Methods (Fig. S2a). The fluorescence of FM4-64 was checked through confocal laser scanning microscopy to visualize the localization of FM4-64-*Xoo*-OMVs in the plant tissue. Roots of 7-day-old Khitish and CR 800 seedlings were incubated in FM4-64-*Xoo-*OMVs and the mock treatment (control). After 30 minutes of incubation in FM4-64-*Xoo-*OMVs, the PM of rice roots showed slight fluorescence, indicating the initiation of fusion of the *Xoo-*OMVs with the rice PM. After 60 minutes or 90 minutes of incubation, the fluorescence in the roots incubated in FM4-64-*Xoo-*OMVs increased considerably, indicating the incorporation of *Xoo-*OMVs into the rice root PM. No fluorescence was observed in the PM of the rice roots incubated in the mock treatment, confirming that the remnant dye was insignificant (Fig. S2c). The fluorescence detection range of FM4-64 is close to that of chloroplasts. Thus, to visualize the membrane of the leaf tissues clearly without misinterpretation due to the auto-fluorescence of the chloroplasts, we used protoplasts isolated from the rice leaf tissues as the system for this experiment. Protoplasts, isolated from the leaves of Khitish and CR 800, were incubated with either FM4-64-*Xoo-*OMVs or the mock treatment. The protoplasts that distinctly showed the PM were chosen for analysis. The ROI was selected from the bright field images on the regions showing the PM (Fig. S2b). It was noticed that like in the roots, the FM4-64-*Xoo-*OMVs were incorporated into the PM of the protoplasts after 60 minutes of incubation (Fig. 4c).

We further checked whether the initial Ca^2+^ signal in the host and the subsequent increase in lipid order of the host membrane are necessary for the incorporation of *Xoo-*OMVs into the rice PM. On chelation of intracellular Ca^2+^ with BAPTA, the insertion of *Xoo-*OMVs in the rice root PM was significantly reduced (Fig. 4a, b). The protoplasts pre-treated with BAPTA manifested a similar decrease in the insertion of *Xoo-*OMVs into the rice PM (Fig. 4c, d). Roots of Khitish and CR 800 seedlings pre-treated with MβCD for 30 minutes displayed no fusion of *Xoo-*OMVs with the rice PM (Fig. 4a, b). Taken together, our data demonstrate that the integration of *Xoo-*OMVs into rice PM is a Ca^2+^-dependent and nanodomain-mediated process.

### Ca^2+^ is crucial for the enrichment of defence-related proteins in the PM of CR 800 after *Xoo*-OMV treatment

The composition of PM proteins is modulated according to the cell’s state, which helps the cell recognise and respond to various abiotic and biotic stimuli (Cao et al., 2016; Fujiwara et al., 2009; Komatsu, 2008). An increase in the abundance of defence-related proteins involved in ABA signalling, ROS bursts, SA signalling, etc., is observed in the rice PM after exposure to Magnaporthe oryzae (Cao et al., 2016) and Xoo (Chen et al., 2007; Fujiwara et al., 2009).

We examined whether Xoo-OMV-induced increase in lipid order mobilises the assembly of specific proteins in the rice PM. To this end, we carried out a quantitative proteomic analysis of membrane proteins from the CR 800 variety, as CR 800 can mount an immune response against Xoo. Specifically, 7-day-old CR 800 seedlings were incubated in Xoo-OMV or sterile water, or pre-treated with EGTA (2 mM) for 30 minutes before incubation in Xoo-OMV for 1 hour. The PM fraction was enriched using the protocol of Uemura et al. (1995). PM proteins were analysed by label-free quantification (LFQ). A total of 3274 proteins were identified; of these, 2689 proteins were present in all sample groups (Control, EGTA-OMV, and OMV), while 54 were exclusively associated with the PM of roots treated with Xoo-OMVs. We found that approximately 430 proteins were upregulated (≥1.5-fold change, p>0.05) in OMV-treated roots, while 351 proteins were upregulated in EGTA-OMV treated roots.

Furthermore, through functional annotations we found that proteins that were upregulated after Xoo-OMV treatment were predominantly involved in cell wall biogenesis, vesicle docking, Ca2+ signalling, and most importantly, in defence against bacteria. None of the above-mentioned signalling pathways were enriched under Ca2+-chelated conditions.

We identified 233 proteins that were significantly upregulated (≥1.5-fold change, p>0.05) in CR 800 roots in response to *Xoo*-OMVs, but not under a Ca^2+^-deprived condition. Through functional annotation analysis of the these differentially abundant proteins (DAPs) using String Database, we found that these proteins are majorly responsible for plant cell wall remodelling and biogenesis (WAK2, WAKL21, CESA6, CESA4, CSLF6, etc.) (Li et al., 2009; Malukani et al., 2020), receptor-like kinases (LRR-RLKs, RLCKs and lectin domain containing kinases etc.) (Mou et al., 2024; Yang et al., 2020), Ca^2+^ signalling proteins (CPK18, CPK17, CML10, CNGC9 etc.) (Bundó and Coca, 2017; Wang et al., 2019), defence response-related proteins (SERK2, NRT1, RBLS2, RGLG2 IRL6, etc.) (Liu et al., 2023), and lipid metabolism proteins (PLD, ACBP2, KCS, etc.) (Kim et al., 2013; Young et al., 1996) (Fig. 5a). Interestingly, we observed that microdomain-associated proteins, like remorins (REM1.4 and REM4.1) (Gui et al., 2016), syntaxins (SYP121 and SYP71) (Bao et al., 2012; Cao et al., 2019), tetraspanins (TET8 and TET11), aquaporins (PIP2-2 and PIP2-5), and annexin are more abundant in OMV-treated roots than in EGTA-OMV-treated roots. Moreover, proteins that positively regulate resistance against *Xoo*, like, HVA22-like proteins, leaf rust resistance protein, RBLS2, RGLG5, etc. and hypersensitive response proteins like PR1a, PR5, callose synthase, dirigent proteins (Paniagua et al., 2017), HIRL2, etc. were also reduced under a Ca^2+^-chelated condition (Fig. 5b; Table 2). Notably, the abundance of immune co-receptors, playing a key role in defence against *Xoo*, like, SERK2, SOBIR1, BAK1 and BRI1 (Chen et al., 2014; Zhang et al., 2021) are elevated in CR 800 PM proteome following treatment with *Xoo*-OMVs in presence of Ca^2+^, but not in Ca^2+^-deprived condition (Fig. 5b). These outcomes are consistent with our previous findings, which demonstrate that *Xoo*-OMV-induced nanodomain enrichment and immune response are impaired in a Ca^2+^-chelated condition.

**Fig. 5.**
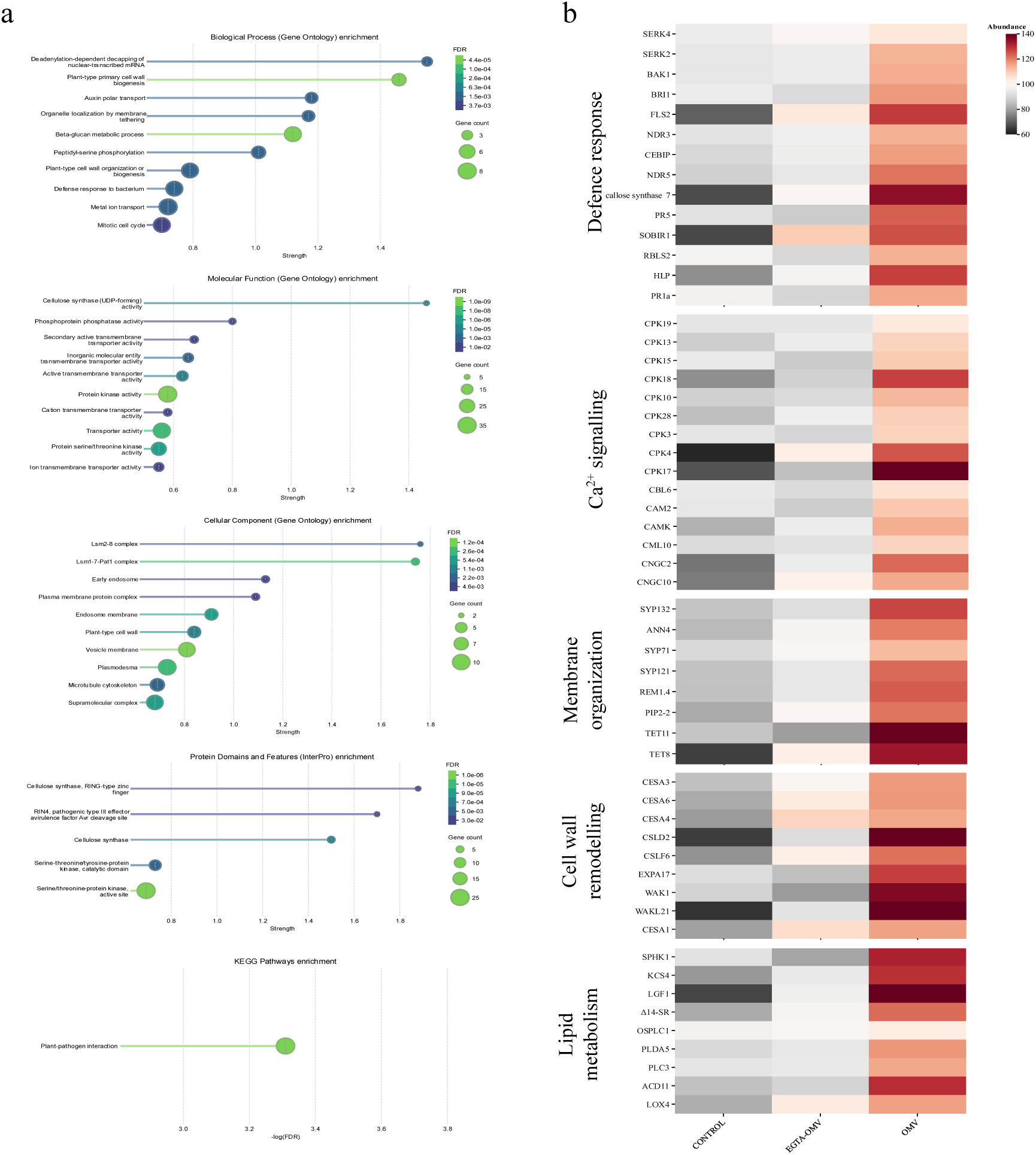
Analysis of the CR 800 PM proteome remodelling after *Xoo*-OMV treatment in native and Ca^2+^-deprived condition. a, Functional annotations of the 233 proteins upregulated (1.5≥Fold-change, p≤0.05) in CR 800 PM after *Xoo*-OMV treatment in normal condition, but not in a Ca^2+^-chelated condition. Enrichment is represented as a lollipop chart created using String Database with strength (log_10_(Expected/Observed)) in the x-axis. b, Heatmap displays the abundance of key proteins involved in important pathways as observed after different treatments, Control (sterile water-1hr), EGTA-OMV (30 min pre-treatment with EGTA (2mM); *Xoo*-OMV-1hr), and OMV (*Xoo*-OMV-1hr), (n=3).

**Table 2:**
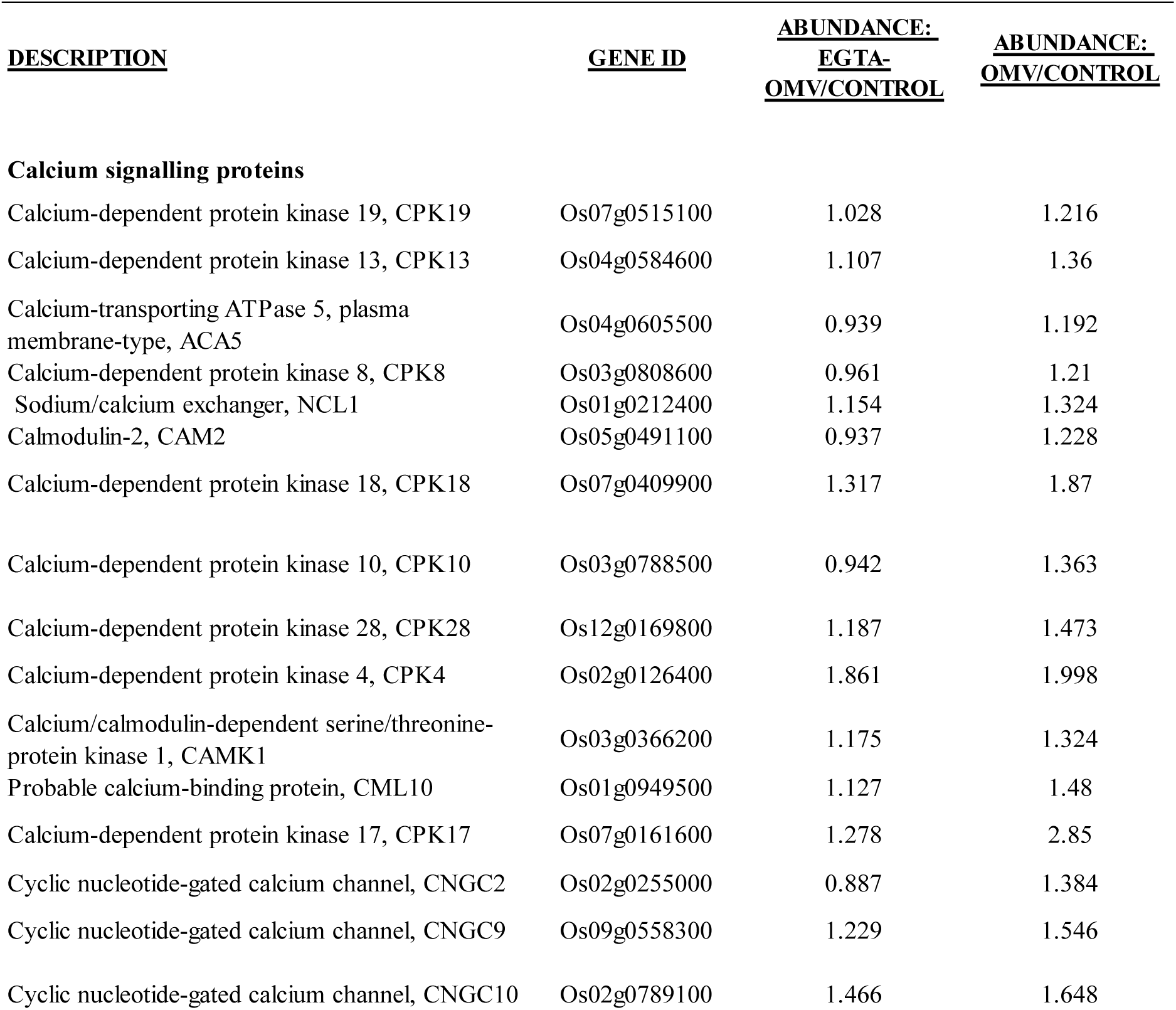
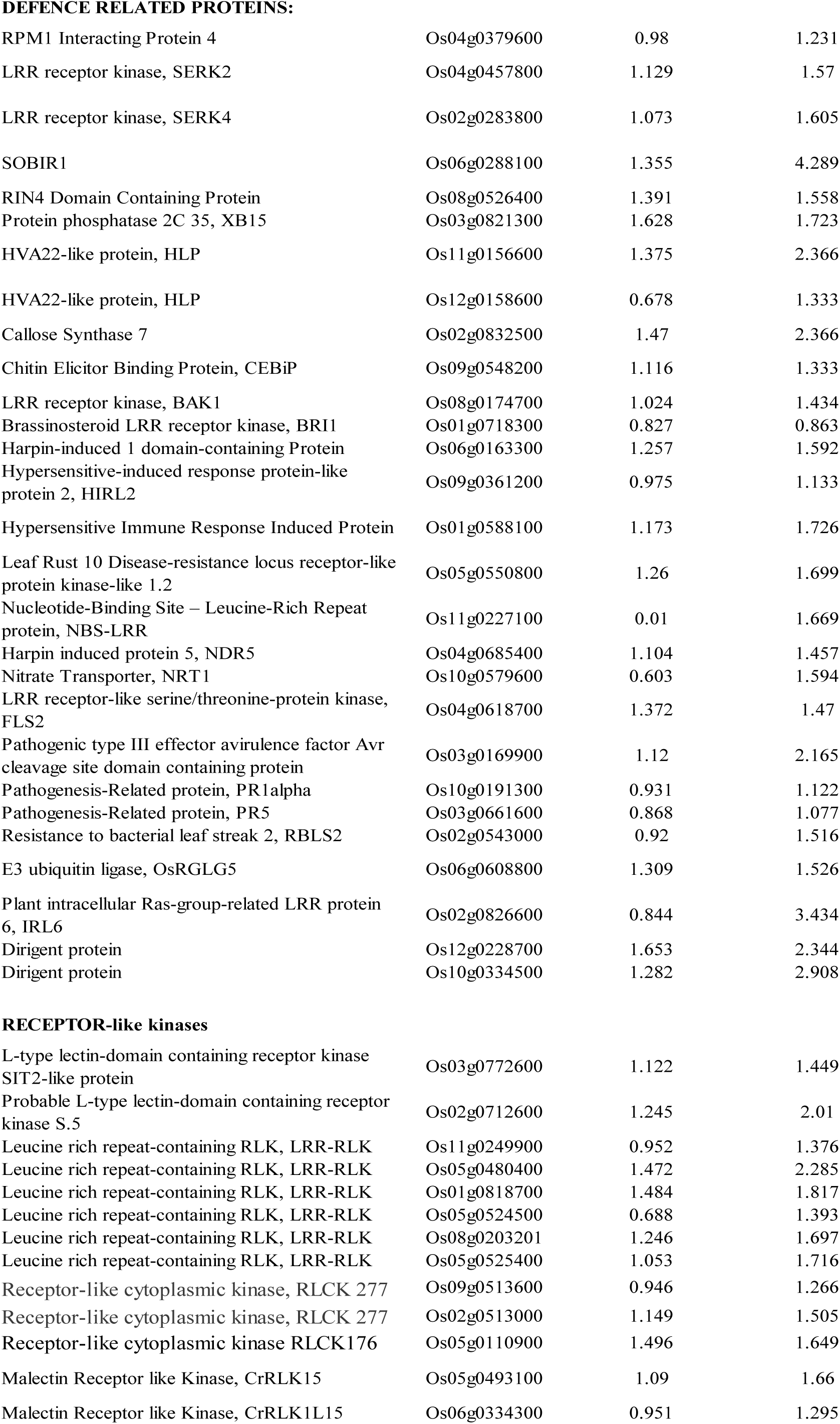
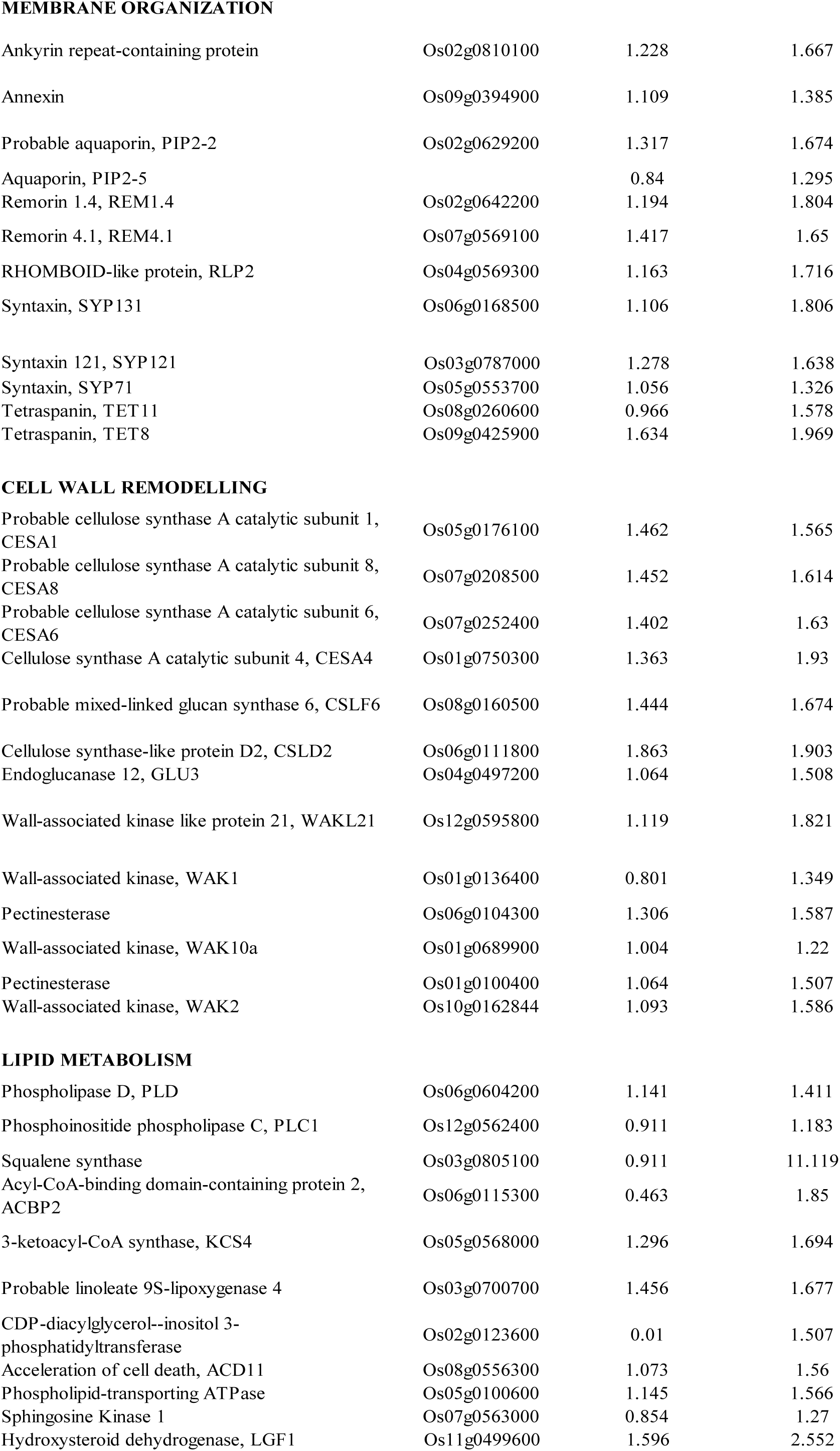
Important proteins found in CR 800 PM with their respective Abundance ratios.

## Discussion

Gram-negative bacteria utilise outer membrane vesicles (OMVs) to enhance their chances of survival within the host, concurrently, the host has evolved to recognise OMVs and mount an immune response against the invading pathogen to protect itself (McMillan and Kuehn, 2021; Schwechheimer and Kuehn, 2015). Emerging evidence depicts the function of phytopathogenic OMVs in plant-pathogen interactions. In Arabidopsis, *Xcc*-OMVs potentiate an immune response via nanodomain-mediated insertion of the OMVs into the plant PM (Bahar et al., 2016; Tran et al., 2022). Additionally, pre-treatment with OMVs, like *Pst*-OMVs in tomato plants or *Xcc*-OMVs in Arabidopsis, can protect the plant from subsequent infection (Chalupowicz et al., 2023; McMillan and Kuehn, 2023). Nevertheless, the molecular mechanism and sequence of events that lead plants to mount an immune response after recognising pathogenic OMVs are still unknown.

Our study aimed to uncover the initial signalling events through which rice perceives *Xoo*-OMVs. We have isolated and categorised the proteins associated with *Xoo*-OMVs to find several MAMPs, virulence-associated proteins, secretory system proteins, DSF-processing enzymes, and hydrolytic enzymes, which indicate the role of *Xoo-*OMVs during invasion and pathogenesis inside the host (Fig. 1, Table 1). We found that several cell wall-degrading enzymes like cellulase, endoglucanase, esterase, etc. (Table 1) are associated with *Xoo*-OMVs, which destabilise the cell wall of the plant to enable the insertion of OMVs in the PM. Meanwhile, cell wall degradation produces DAMPs that can alert the host. Performing quantitative proteome analysis of CR 800 roots, we observed an increase in the abundance of chitin elicitor binding protein (CEBiP) that recognises the cell wall-derived oligosaccharides as DAMPs (Fig. 5b) to mount an immune response (Yang et al., 2021). Interestingly, *Xoo*-OMV-treated rice proteome shows increased abundance of (i) cell wall-remodelling enzymes (endoglucanase, pectinesterase, WAKs) to produce more DAMPs and amplify the immune response and modify the cell wall, and (ii) cell wall biosynthesis enzymes (CESAs, CSFL6) to repair the damaged cell wall. *Xoo*-OMVs contain iron and phosphate channels (Table 1) to ensure optimum virulence (Chatterjee et al., 2003; Subramoni et al., 2012). We also observed an increase in abundance of several metal ion transporters in rice root PM after *Xoo*-OMV treatment, suggesting a probable competition for available nutrients between the plant and the pathogen (Fig. 5a). The proteins carried by the *Xoo-*OMVs reflect that OMVs consist of numerous proteins that not only enhance virulence of *Xoo* but also defend the bacteria against plant’s immune responses and promote the chance of survival inside the host. At the same time, the plant recognizes the OMVs and responds by recruiting necessary proteins in its PM.

MAMPs, such as flg22 and elf18, are known to induce Ca^2+^ signals in Arabidopsis (Ranf et al., 2014). We have observed that, *Xoo*-OMVs which carry a complex cocktail of MAMPs, induces a gradual Ca^2+^ signal in the elongation zone of the rice root after 10 minutes of *Xoo*-OMV application. Live cell Ca^2+^ imaging with Arabidopsis Col-0-RGECO1 seedlings revealed that the spatiotemporal resolution of the signal is unique compared to other individual MAMPs (Fig. 2). OMVs consist of lipopolysaccharides and MAMPs like flg22, EF-Tu, extracellular peptidase, cellulase, and other extracellular secretion system proteins (Table S1), the Ca^2+^ signals that we have observed is not specific to any particular MAMP; rather, it is a response of the root cells towards a mixture of MAMPs. Our results demonstrate that this rapid Ca^2+^ signal is the key to *Xoo*-OMV perception by rice, and alongside ROS serves as another hallmark of OMV perception by plants.

It is well-established that plant pathogen-derived OMVs can trigger an immune response in the host plant (Bahar et al., 2016; Chalupowicz et al., 2023; McMillan et al., 2021). CR 800 (expressing *Xa21, xa3*, and *xa15*) can display a significant immune response against *Xoo*-OMVs, while Khitish (expressing no R genes) is susceptible to *Xoo* infection and cannot mount any immune response against *Xoo*-OMVs. These results are consistent with previous reports (Liu et al., 2020; Thomas et al., 2016), where the resistant rice line, expressing *Xa21,* shows elevation in the expression of *PR10b* while the susceptible rice line does not show elevated *PR10b* expression. Essentially, we noted that blocking this early Ca^2+^ signal by Ca^2+^-chelation results in a weakened immune response in the resistant variety CR 800 (Fig. 2). Through further investigation, we have observed that *Xoo*-OMV-induced nanodomain enrichment is also impaired in Ca^2+^-chelated roots (Fig. 3). As ordered PM is necessary for OMV insertion (Tran et al., 2022), the disordered membrane of EGTA-treated roots show reduced *Xoo*-OMV insertion (Fig. 4). Temporal analysis of the events elucidates, Ca^2+^ signalling as one of the earliest event during *Xoo*-OMV perception (10 minutes after application), followed by nanodomain enrichment (30 minutes after *Xoo*-OMV application) and insertion of OMVs into the rice PM (after 60 minutes of incubation in *Xoo*-OMVs). Tran et al., 2022 showed that *Xcc*-OMVs get inserted in the Arabidopsis PM within 10 minutes. The delay in the incorporation of the *Xoo-*OMVs in rice PM (60 minutes) is probably due to the differential composition of the cell walls of rice and Arabidopsis, as rice roots are more densely packed and rigid than Arabidopsis roots (Vogel, 2008).

Based on these observations, we hypothesized that Ca^2+^ signalling led to aggregation of specific proteins in the rice PM, which shapes the response of rice towards *Xoo*-OMVs. To verify our hypothesis, we performed quantitative proteomics analysis of CR 800 root PM after *Xoo*-OMV application, both in the presence and absence of Ca^2+^-chelator (EGTA). We observed that several defence response-related proteins were significantly upregulated in CR 800 OMV-treated roots but not in EGTA-OMV-treated roots, validating the compromised immune response displayed on Ca^2+^-chelation. Several sterol and lipid biosynthesis pathway-related proteins were also more abundant after *Xoo*-OMV application, which leads to a highly ordered PM (Fig. 5, Table 1). Contrastingly, in a Ca^2+^-deprived condition, these proteins were not significantly upregulated, which might lead to a disordered membrane, as manifested in Di-4-staining assay (Fig. 3). Similarly, membrane organisation and nanodomain-associated proteins like remorins, syntaxins, annexin, tetraspanins, etc, which aid in vesicle fusion and docking, are highly abundant in OMV-treated roots, but not in roots treated with EGTA-OMV-treated roots (Fig. 5, Table 1).

Taken together, our outcomes reveal that *Xoo* releases OMVs that comprise abundant MAMPs and virulence-associated proteins. These MAMP-loaded *Xoo*-OMVs are sensed by the plant, via a Ca^2+^ signal generation. The Ca^2+^ signal enables the organization of a nanodomain-enriched membrane possibly by upregulating the sterol biosynthesis pathway and leads to the aggregation of specialized defence proteins in the rice PM. These proteins transduce the signal downstream and generate an immune response against the pathogen. Remorins, annexins, ankyrin repeat-containing proteins, and SNARE proteins like syntaxins, aggregate in the membrane, which might aid in the insertion of *Xoo*-OMVs into the rice PM. Such nanodomain congregation is necessary to prepare the cell for a subsequent response to combat the pathogenic challenge by recruiting the defence-related proteins into signalling hubs. In a Ca^2+^-deprived condition, these proteins are not efficiently assembled in the PM, resulting in a perturbed recognition of *Xoo*-OMVs by the plant and a disrupted immune response. Although, Khitish, the susceptible variety, could perceive the *Xoo-*OMVs through Ca^2+^ signal generation, nanodomain enrichment and insertion of *Xoo*-OMVs into the rice PM, it failed to evoke any immune response due to the absence of R genes and the associated repertoire of proteins (Figure 6).

**Fig. 6.**
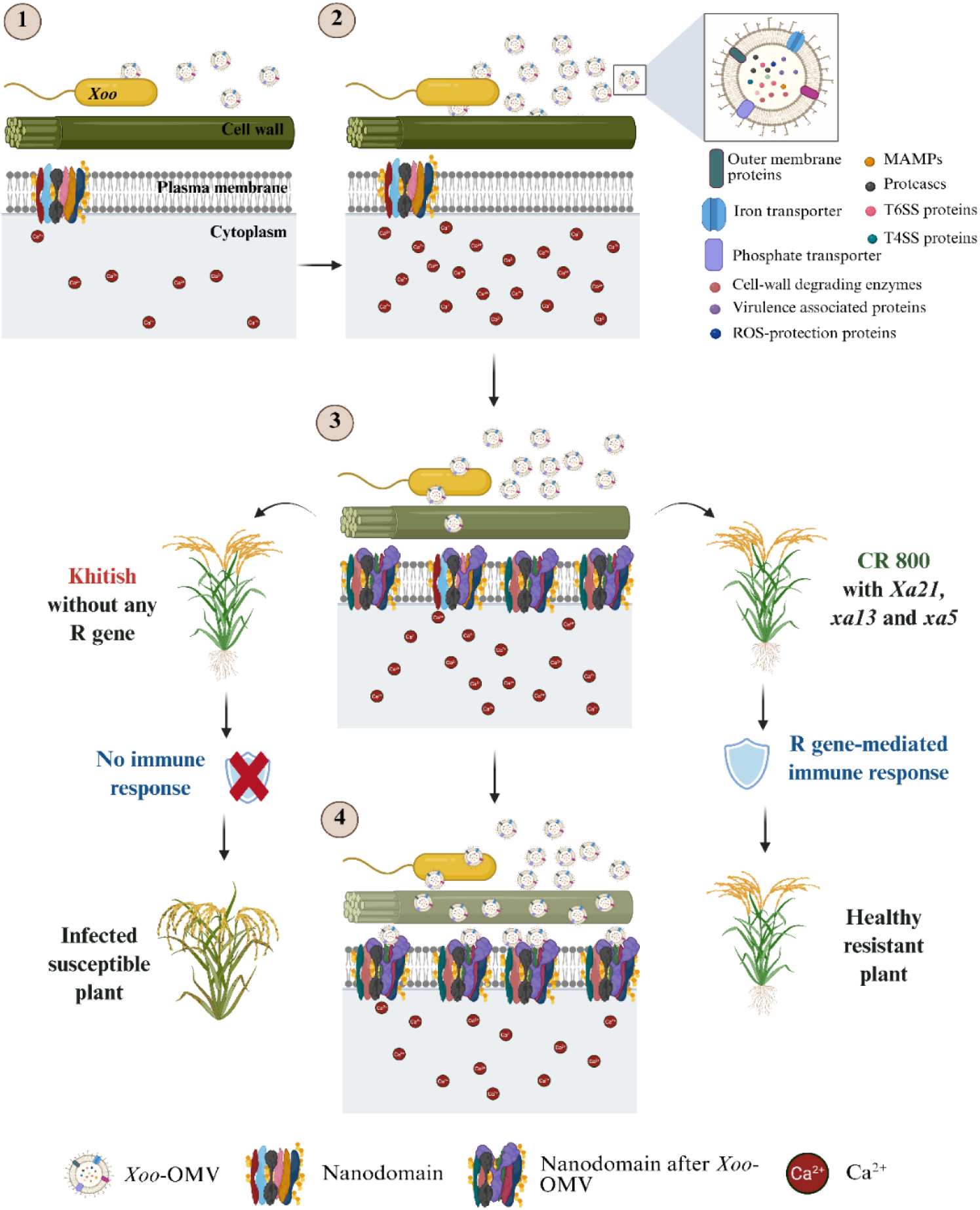
Ca^2+-^dependent perception of *Xoo*-OMVs by rice via nanodomain-mediated PM proteome reorganization. *Xoo*-OMVs are associated with MAMPs and virulence-associated proteins. Through these MAMPs, the rice cell recognizes the *Xoo*-OMVs by evoking a Ca^2+^ signal. This leads to nanodomain enrichment in the rice PM and shifts the PM proteome towards the defence response pathway. It also enables insertion of *Xoo*-OMVs into the rice PM. Both Khitish and CR 800 perceive the *Xoo*-OMVs similarly, but, the resistant variety CR 800, expressing R genes (*Xa21*, *xa13, xa5*) mounts an immune response against the pathogen, while Khitish the susceptible variety without any R gene cannot evoke any immune response. If the early Ca^2+^ signal is blocked, *Xoo*-OMV perception and following immune response are impaired in CR 800.

Remarkably, specific nanodomains are organized with appropriate proteins and function in a signal-specific manner (Bücherl et al., 2017). However, the mechanism of such signal-specific nanodomain assembly is still elusive. We propose that the precise and spatiotemporally unique Ca^2+^ signals exhibited in response to different biotic and abiotic stimuli can stipulate such a signal-specific nanodomain formation. Further research is required to uncover the molecular mechanism of Ca^2+^-mediated nanodomain configuration to understand the recruitment of the various signalling components at the plant plasma membrane.

## Acknowledgments

This work was funded by the SRG/2022/002293 grant from the Department of Science and Technology Government of India. We thank Dr. H. K. Patel from CSIR-CCMB for providing us with the *Xoo* strains that have been used in the study. We are thankful to the Rice Research Station, Chinsurah, West Bengal, for providing us with the seeds of Khitish and CR 800. We acknowledge the Central Instrumentation Facility (CIF) of CSIR-Indian Institute of Chemical Biology (CSIR-IICB). We are thankful to DBT-ILS for the TEM and ESI-MS/MS experiments. We Acknowledge Mr. Sandip Barman for critically reading the manuscript for editing.

## Author contributions

SB conceptualized the project and arranged the funding. IM and SB designed the experiments. IM and HD performed the experiments. IM and SB analysed the data. IM and SB wrote the manuscript.

## Data availability

All the data discussed in the manuscript are included as supplementary information. All of the proteins identified from the *Xoo*-OMV-associated proteome and CR 800 PM-associated proteome analysis are listed in the supplementary tables, Table 1 and Table 2, respectively. Raw files of proteomic analysis are available on request.

## Supporting Information

The following Supporting Information is available for this article:

**Fig. S1** Temporal nanodomain enrichment in rice roots after exposure to *Xoo-*OMVs in Ca^2+^-chelated condition.

**Fig. S2** Insertion of *Xoo-*OMVs into the PM of rice with respect to time.

**Fig. S3** Overview of CR 800 PM proteome.

**Fig. S4** Functional annotation of upregulated proteins.

**Table S1** All the proteins identified in the Xoo-OMVs.

**Table S2** All the proteins identified in the CR 800 PM.

**Video S1** Ca^2+^ signal in Arabidopsis Col-0-RGECO1 root after Xoo-OMV application.

