## Supplemental files for "Ca^2+^-driven nanodomain enrichment and plasma membrane proteome remodelling enable bacterial outer membrane vesicle perception in rice"

**Fig. S1** Temporal nanodomain enrichment in rice roots after exposure to *Xoo*-OMVs in Ca$^{2+}$-chelated condition. a, and b, Di-4-ANEPPDHQ staining of Khitish and CR 800 root respectively, after 30, 60, and 90 minutes of *Xoo*-OMV application, with or without pre-treatment, as mentioned. c, the GP values of the Khitish and CR 800 PM respectively at each 30, 60 and 90 minutes are presented in a box plot. d and e, the GP values of Khitish and CR 800 root PM, at 60 and 90 minutes, respectively, after Xoo-treatment, with or without Ca$^{2+}$-chelation, are plotted as box plots. All box plots are created using GraphPad Prism software, and statistical significance is calculated by One-way ANOVA with p-value was taken as 0.05 > ns, not significant, *$P \leq 0.05$; **$P \leq 0.01$, ***$P \leq 0.001$, ****$P \leq 0.0001$.

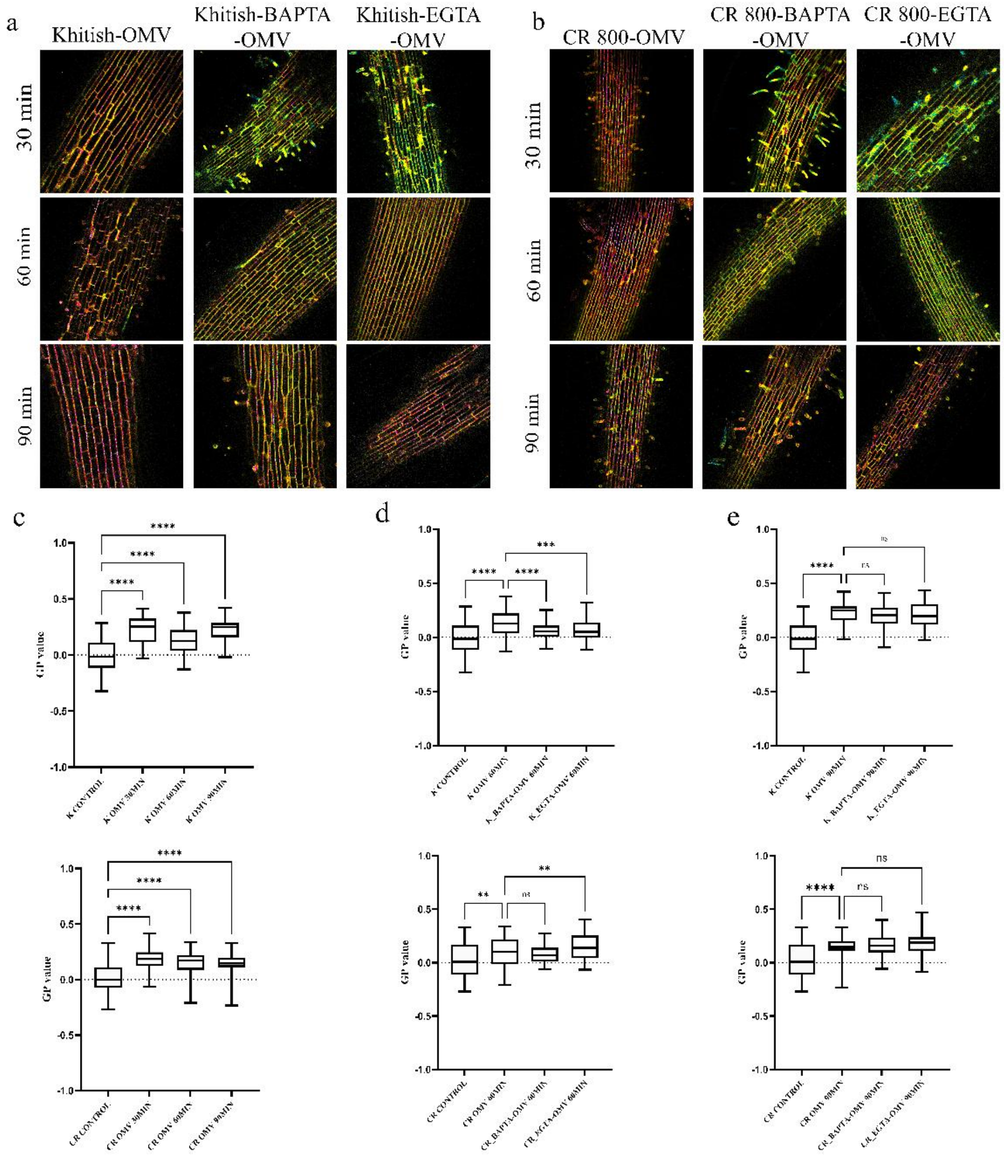

**Fig. S2** Insertion of *Xoo*-OMVs into the PM of rice with respect to time. a, schematic representation of protocol of Xoo-OMV staining with FM4-64. b, ROI selection in rice protoplast for analysis. c, insertion of Xoo-OMVs in rice PM with respect to time. d, the histograms showing relative fluorescence units at each time point, were analysed using GraphPad Prism software by One-way ANOVA with p-value was taken as 0.05 > ns, not significant, *P ≤ 0.05; **P ≤ 0.01, ***P ≤ 0.001, ****P ≤ 0.0001.

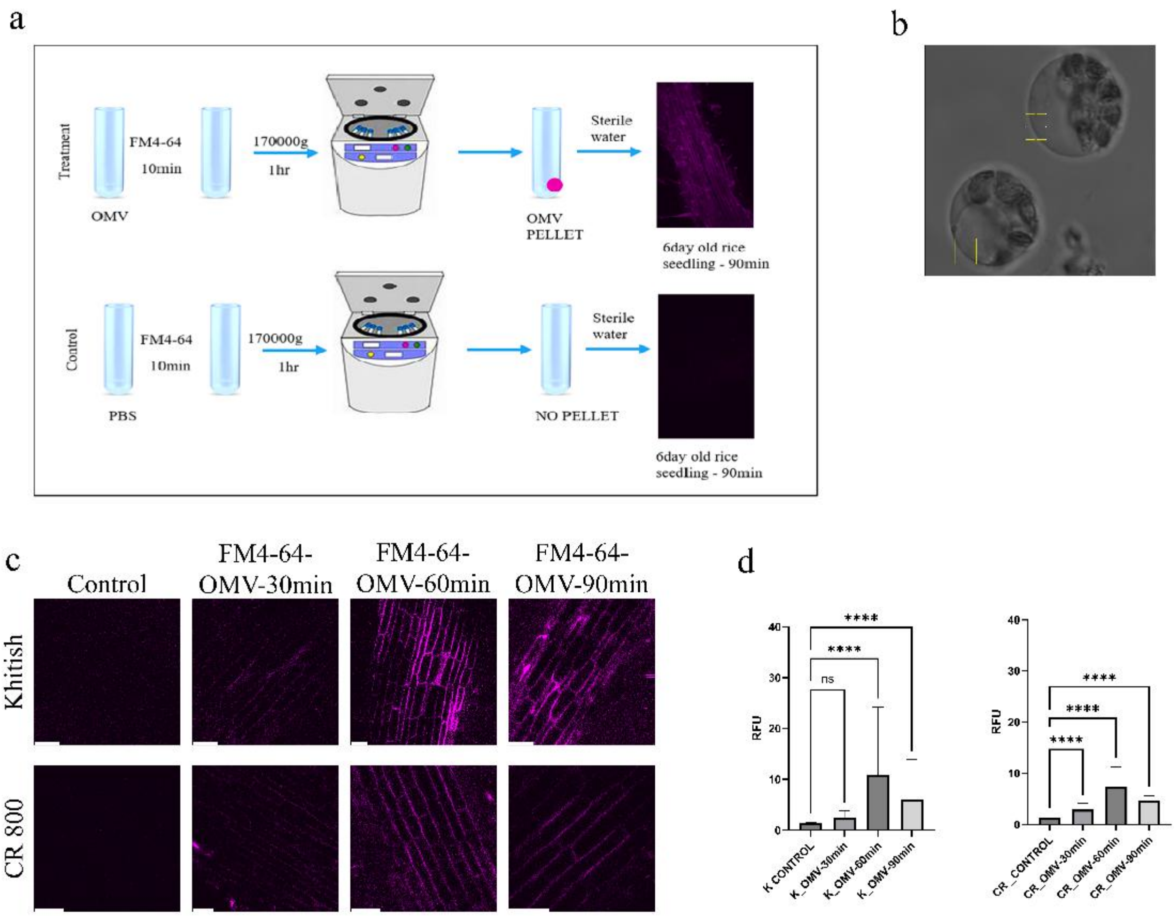

**Fig. S3** Overview of CR 800 PM proteome. a, Heatmap of all the 3274 proteins identified in the CR 800 PM preparation, based on Grouped Abundance of each protein in each treatment (n=3). b, PCA analysis of the samples Control (blue), EGTA-OMV (orange), and OMV (green), shows that PM proteome shifts distinctly after *Xoo*-OMV treatment, but this shift is perturbed in a $Ca^{2+}$-chelated condition. c, volcano plot depicting the proteins differentially abundant in Xoo-OMV treated sample, with respect to the control sample. Significantly upregulated proteins are highlighted in red and significantly downregulated protein are shown in green. d, Venn diagram demonstrating the presence of different proteins in different sample groups. All of the above plots are generated through Thermo Proteome Discoverer software.

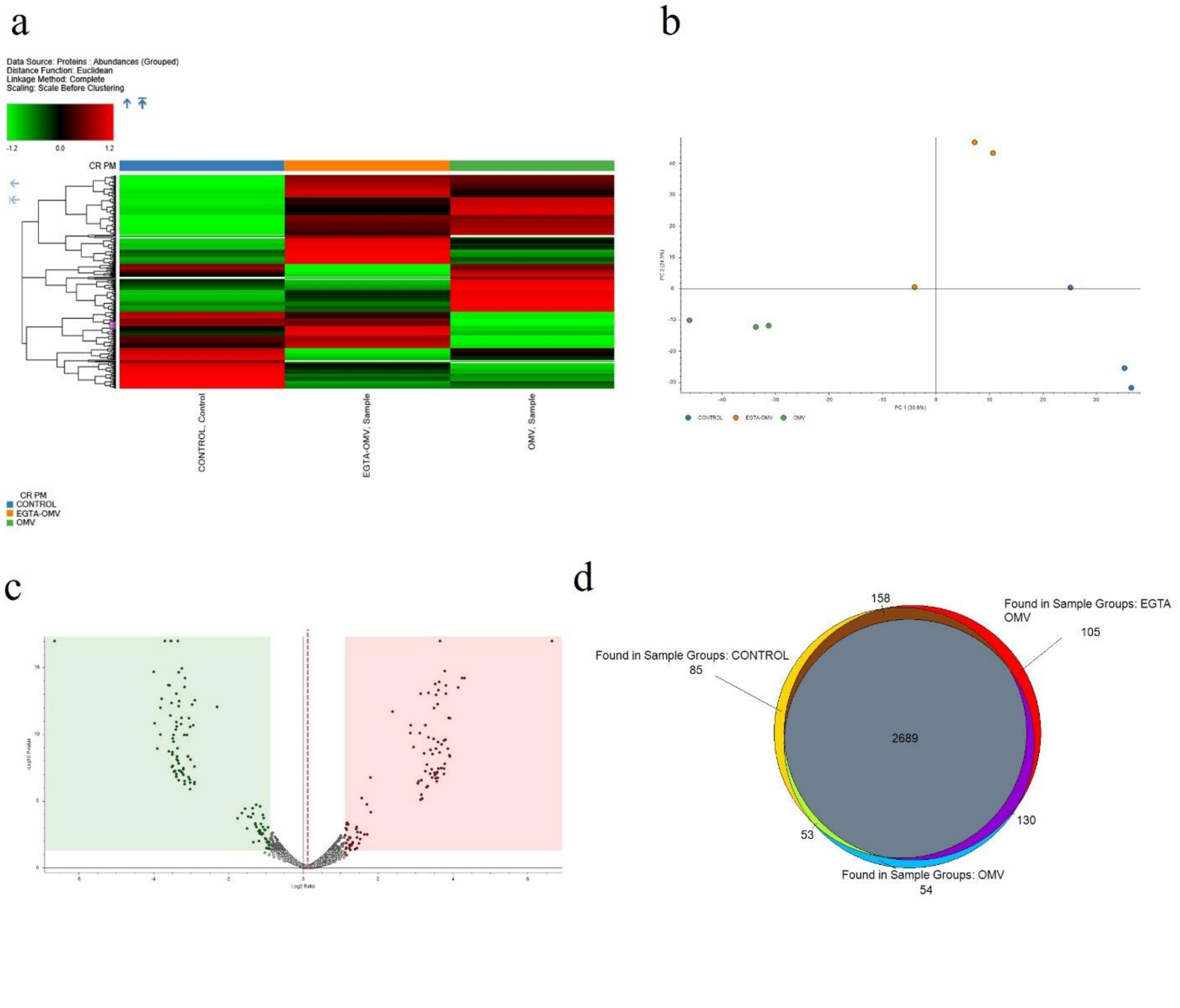

**Fig. S4** Functional annotation of upregulated proteins. a, functional annotations of 430 proteins significantly upregulated in CR 800 PM after 1 hour of Xoo-OMV treatment. b, functional annotations of 351 protein significantly upregulated in CR 800 PM after 30 min EGTA pre-treatment and 1 hour of Xoo-OMV treatment.

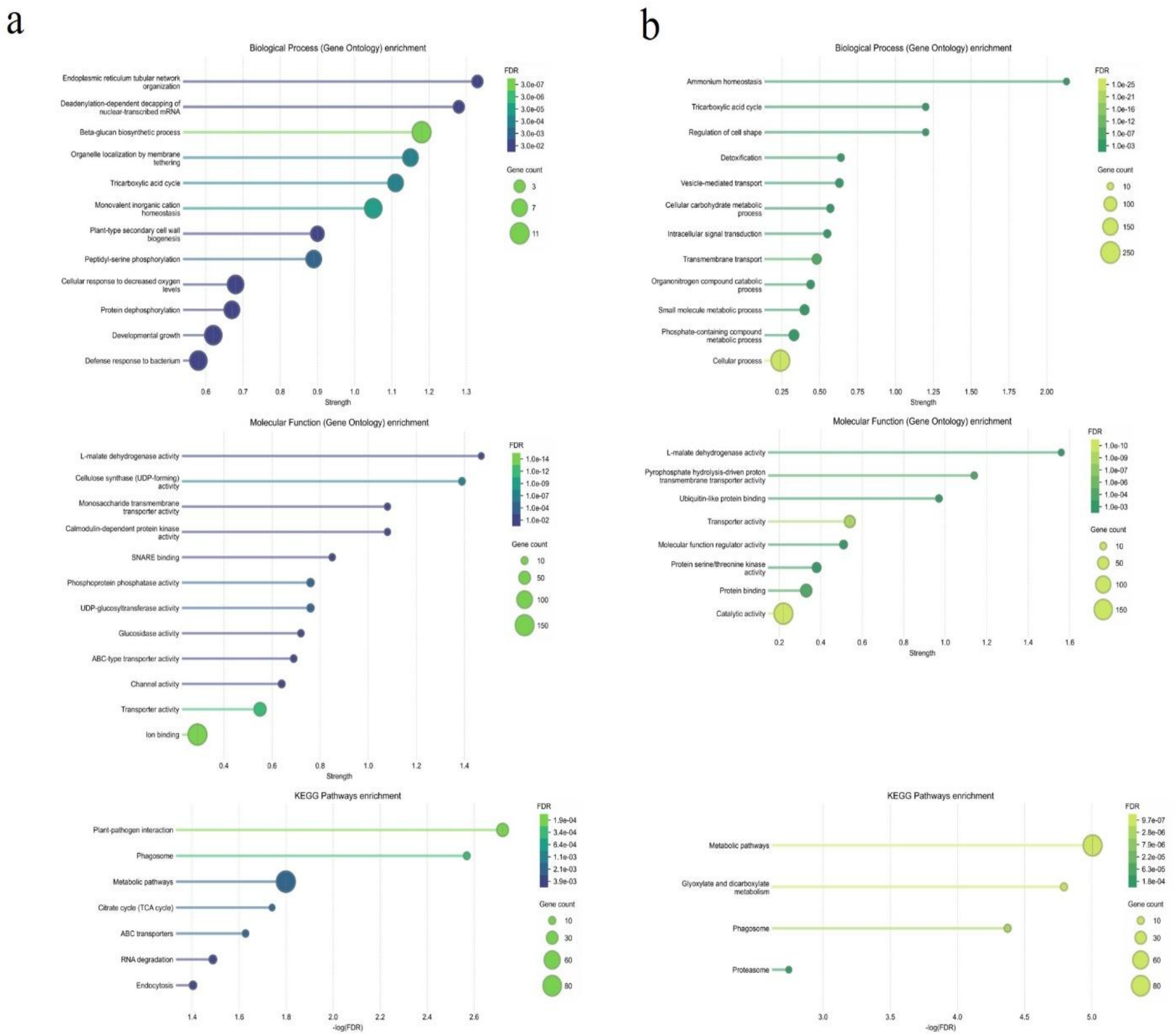

**Table S1** All the proteins identified through proteomics of purified Xoo-OMVs, showing at least 2 unique peptides, filtered through Percolator.
**Table S2** All the proteins identified in CR 800 PM using Sequest HT along with Percolator.

**Video/Movie S1** Unique Ca2+ signal is observed in the root elongation region of 4-day-old Arabidopsis Col-0-RGECO1 seedings treated with Xoo-OMV after 5 minutes of imaging.
